# P-cadherin and desmoglein-2 interact as strand-swap dimers and facilitate desmosome assembly

**DOI:** 10.1101/2025.09.15.676363

**Authors:** Shipeng Xu, Nishadi Mudiyanselage, Sanjeevi Sivasankar

## Abstract

Desmosomes are essential adhesive junctions that mediate the integrity of tension prone tissue like the heart and skin. The classical cadherin, P-cadherin (Pcad), is known to play an important role in forming desmosomes. However, the molecular mechanisms by which Pcads enable desmosome assembly are unknown. Here, we combine single molecule Atomic Force Microscopy, super-resolution and confocal imaging, mutagenesis assays, and atomistic simulations to resolve the structural mechanisms by which Pcads facilitate the formation of desmosomes. We show that Pcads interact with the desmosomal cadherin Desmoglien-2 (Dsg2) on opposing cells, by mutually swapping β-strands terminated by conserved Tryptophan residues. A key determinant of this trans heterophilic strand-swap dimerization is the flexibility of a hinge on the swapped β -strands: stiffening this hinge reduces the likelihood of heterophilic strand-swap dimer formation. We demonstrate that while desmosome formation is impaired in cells that either lack classical cadherins or that express strand-swap deficient Pcad, introduction of strand-swap competent Pcad into these cells rescues desmosome assembly. We show that the heterophilic dimers formed by Pcad and Dsg2 interact in a robust and persistent manner, which retains Pcad within the desmosome throughout the maturation process.

**Significance:** Desmosomes are essential intercellular junctions that mediate the integrity of tissue like the heart and skin. Desmosomal defects are associated with numerous skin disorders, heart diseases, and cancer. The adhesive protein P-cadherin is known to play an important role in forming desmosomes. However, the molecular mechanisms by which P-cadherins enable desmosome assembly are unknown. Here, we show that desmosome formation is facilitated by the direct interaction of P-cadherin with the transmembrane desmosomal adhesion protein Desmoglein-2. We demonstrate that P-cadherin and Desmoglien-2 interact robustly by swapping flexible portions of their protein backbone and that this molecular complex nucleates desmosome formation. Our results resolve the biophysical basis by which different adhesive proteins interact and mediate stable desmosome assembly.

## Introduction

Multicellular organisms rely on a range of cell-cell adhesion junctions to mediate tissue morphogenesis and maintain tissue integrity ^1^. A key adhesion protein in these junctions are cadherins, a superfamily of calcium-dependent transmembrane proteins that mediate homophilic and heterophilic interactions between adjacent cells. There are two major cadherin-based cell-cell junctions: Adherens Junctions (AJs), which are which are present in all soft tissue, and are composed of classical cadherins, and Desmosomes, which are present in tissue such as the skin and heart, and are composed of desmosomal cadherins ^2,3^. AJs and desmosomes are not functionally isolated: accumulating evidence suggests that AJs are critical for the initiation of desmosome assembly and for its subsequent maturation ^4,5^. Despite the functional interdependence between AJs and desmosomes, the molecular details of heterophilic interactions between their components are not completely understood.

There are two types of desmosomal cadherins: desmocollins (Dsc) comprised of three isoforms and desmogleins (Dsg) comprised of four isoforms. Previous studies have implicated two classical cadherins - E-cadherin (Ecad) and P-cadherin (Pcad) - in desmosome assembly. These studies show that Ecad and Pcad transiently colocalize with desmosomal cadherins such as isoform 2 of Dsg (Dsg2) and form mixed junctional complexes ^6^. As desmosome mature, classical cadherins segregate to AJs, while Dsg2 clusters within raft-associated membrane domains. This suggests that temporally regulated heterophilic interactions between Dsg2 and classical cadherins support initiation and remodeling of desmosomes ^7,8^.

Blocking antibodies against both Ecad and Pcad have been shown to not only prevent AJ formation but also disrupt desmosome assembly ^9^. Additionally, Pcad has been shown to maintain desmosomal integrity and adhesive function under conditions of Dsg3 insufficiency, suggesting that Pcad interacts with other desmosomal cadherins ^2,10^. Early electron microscopy studies identified Ecad at nascent desmosomal sites ^11^, and later work showed that Ecad directly interacts with Dsg2 to facilitate early desmosome formation ^8^. However, it is unknown if Pcad also directly interacts with desmosomal cadherins such as Dsg2.

The binding interfaces on Ecad and Pcad share several similarities. Both classical cadherins interact via two major interfaces: a cis interface that facilitates binding between cadherins on the same membrane and a trans interface that enables interactions between cadherins on opposing membranes (**Fig. 1a**). Cis interactions occur via the interaction of a conserved isoleucine (I175) residue on one cadherin and a valine (V81) on an adjacent cadherin, and promote cadherin clustering ^12^. In contrast, trans interactions occur between opposing cadherins, and the most robust trans binding conformation is the strand-swap dimer (S-dimer), which is formed by the exchange of conserved tryptophan (W2) residues on extracellular 1 (EC1) domain between binding partners ^13–16^.

**Figure 1.**
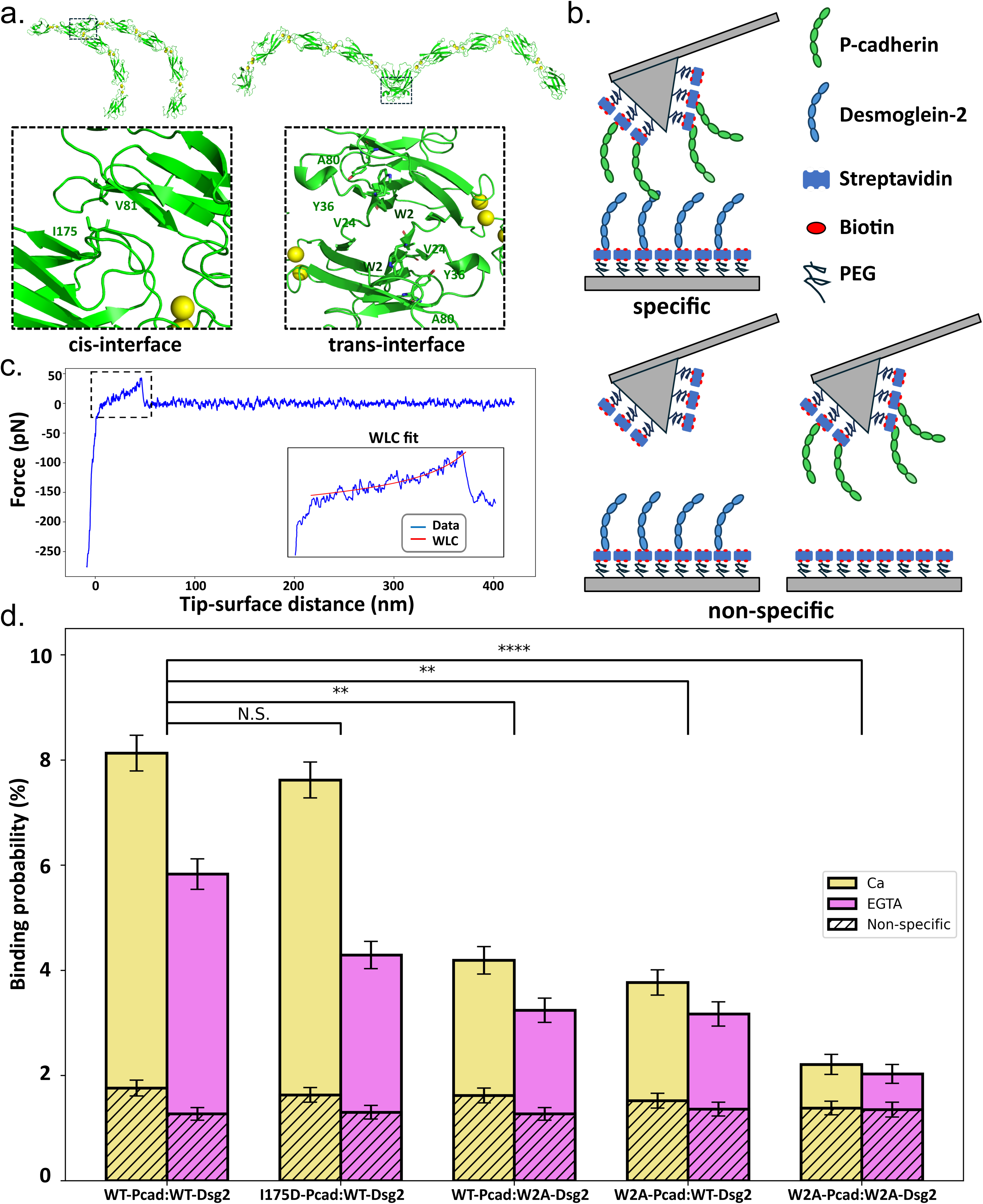
P-cadherin binds to Desmoglien-2 through trans interface. **(a)** Pcad homophilic binding conformations. Left panel: cis interactions between Pcads are mediated through hydrophobic interactions between Val81 and Ile175. Right panel: trans strand-swapping interactions between Pcads occur via the exchange of Trp2 residues between opposing cadherins. **(b)** Schematic representation of the single-molecule AFM binding assay. Upper panel: scheme used to evaluate Pcad–Dsg2 heterophilic interactions. Full ectodomains of Pcads and Dsg2s were immobilized on the functionalized surfaces of the CS and the AFM tip. Lower panels: scheme of control experiments assessing non-specific interactions. Binding probability was measured between a CS substrate decorated with cadherin proteins and an AFM tip lacking cadherins (left), as well as between a protein-free substrate and an AFM tip coated with cadherins (right). **(c)** Representative force–distance curve illustrating a specific interaction event. The PEG tether extension was fitted to the WLC model (red line) in the zoomed-in view. **(d)** Specific binding probabilities between various combinations of Pcad, Dsg2, and their mutants measured under Ca²⁺ (yellow) or EGTA (magenta) conditions, obtained from three independent replicates. Non-specific binding probabilities (hatched black) were obtained from four independent replicates. Binding probability between WT-Pcad and WT-Dsg2 was comparable to I175D-Pcad and WT-Dsg2. Interactions between WT-Pcad and W2A-Dsg2 or W2A-Pcad and WT-Dsg2 showed significantly reduced binding. The interaction between W2A-Pcad and W2A-Dsg2 indicated further reduced binding to a level comparable to non-specific control. For all specific measurements, n = 6075. For all non-specific measurements, n = 4050. All statistical analyses were performed in comparison with WT-Pcad:WT-Dsg2 group. For I175D-Pcad:Dsg2, the P-value is 0.407. For WT-Pcad:W2A-Dsg2, the P-value is 1.05E-3. For W2A-Pcad:WT-Dsg2, the P-value is 4.44E-3. For W2A-Pcad:W2A-Dsg2, the P-value is 1.89E-5. Error bars represent standard error calculated using bootstrapping with replacement.

Unlike classical cadherins which largely interact homophilically, desmosomal cadherins largely exhibit a heterophilic binding paradigm in which Dsgs form strand-swapped heterodimers with Dscs by swapping W2 residues ^17^. This pairing preference is likely dictated by electrostatic complementarity between opposing EC1 interfaces and mirrors the structural architecture of the S-dimer observed in classical cadherins ^17^. Given the structural similarities between classical and desmosomal cadherins, analyzing heterophilic interactions between Pcad and Dsg2 within a ‘cis-dimer’ or ‘S-dimer’ framework provides mechanistic insights into interactions between classical and desmosomal cadherins.

To investigate the molecular interaction between Pcad and Dsg2, we therefore engineered specific mutants that selectively disrupt either the cis-dimerization on Pcad or S-dimer formation on Pcad and Dsg2. We analyzed the interactions of WT and mutant Pcad and Dsg2 using single molecule atomic force microscopy (AFM) and atomistic computer simulations. We also evaluated Pcad expression and localization dynamics during desmosome assembly using confocal and super-resolution microscopy. Our data reveal that unlike Ecad which forms Ca²⁺-independent cis dimers with Dsg2 ^8^, Pcad forms a Ca²⁺-dependent heterophilic S-dimers with Dsg2. Furthermore, unlike Ecad which transiently associates only with nascent desmosomes ^8^, Pcad stably associates with the demosome throughout the assembly process. We show that a critical determinant for the formation of heterophilic S-dimers between Pcad and Dsg2 is the flexibility of the β-strand on Pcad which anchors the swapped W2 residue. Our results resolve the biophysical basis by which classical cadherins interact with desmosomal cadherins and mediate stable desmosome formation.

## Results

### Pcad and Dsg2 form strand-swap dimers

We characterized the interaction between Pcad and Dsg2 using single-molecule AFM binding assays. Wild-type (WT) and mutant forms of the complete extracellular regions (EC1–5) of Pcad and Dsg2 were expressed in suspension-adapted mammalian cells and site-specifically biotinylated at their C-termini. The purified proteins were then immobilized on glass coverslips (CS) and AFM cantilever tips via polyethylene glycol (PEG) tethers (**Fig. 1b**). During each AFM experiment, the protein-coated cantilever tip was brought into proximity with the protein-coated coverslip surface to enable cadherin-mediated interactions. Retraction of the cantilever from the surface generated force-extension curves, which were recorded (**Fig. 1c**). To isolate single-molecule unbinding events and exclude non-specific interactions, we fit the stretching of the PEG tether to a worm-like chain (WLC) model (**Fig. 1c**). The frequency of specific unbinding events was quantified and summarized across experiments.

We first measured the binding probability of WT-Pcad and WT-Dsg2. Our measurements showed a direct, Ca²⁺ dependent interaction between Pcad and Dsg2 ectodomains (binding probability of 8.0% in Ca²⁺ and 5.7% in the Ca²⁺ chelator EGTA; **Fig. 1d**). Given previous findings that Ecad interacts with Dsg2 via its cis-interface and that Pcad shares a homologous hydrophobic residue implicated in this interaction ^8^, we hypothesized that Pcad may also bind Dsg2 in a cis-orientation. To test this, we abolished Pcad cis binding by mutating the conserved Ile175 residue to a negatively charged Asp (I175D). Surprisingly, the binding probability of the I175D-Pcad and WT-Dsg2 was very similar to WT-Pcad and WT-Dsg2 suggesting that this interaction does not rely on the cis-interface of Pcad (7.8% in Ca²⁺ and 4.5% in EGTA; **Fig. 1d**). Next, to determine whether Pcad interacts with Dsg2 via its S-dimer interface, we mutated the swapped W2 in Pcad to an Ala (W2A) and measured the interaction of W2A-Pcad with WT-Dsg2. Simultaneously, we also generated a W2A-Dsg2 mutant and measured its interaction with WT-Pcad. Our measurements showed a significantly reduced interaction probability between both W2A-Pcad:WT-Dsg2 and WT-Pcad:W2A-Dsg2 (4.0% in Ca²⁺ and 3.0% in EGTA; **Fig. 1d**). When we measured the interaction of W2A-Pcad and W2A-Dsg2, the binding probability dropped to background levels (2.0% in both Ca²⁺ and EGTA; **Fig. 1d**). Together, these findings suggest that Pcad–Dsg2 heterophilic interactions occur via a trans strand-swap mechanism involving the swapping of W2 residues from both molecules.

To validate the strand-swap interaction in a cellular context, we performed co-immunoprecipitation (co-IP) assays in Ecad and Pcad knock out A431 cells (Ecad/Pcad KO) ^18^ that we rescued with either WT or mutant Pcad tagged with mCherry at the C-terminus. Given that Ca²⁺ dependent classical cadherins initiate desmosome assembly ^4,9^, prior to performing the co-IP we used a ‘calcium-switch’ protocol to uniformly trigger desmosome assembly in all samples. In our protocol, a confluent monolayer of rescued A431 cells was subjected to calcium depletion using Ca²⁺-free culture medium. This treatment caused the cells to round up and dissociate. Calcium was then reintroduced by switching to a high-Ca²⁺ medium to promote reestablishment of intercellular junctions and initiate desmosome assembly. After overnight high-Ca²⁺ treatment, cells were lysed. The lysates were immunoprecipitated with anti-mCherry antibody to isolate Pcad and associated binding partners. Western blot analysis demonstrated that while Dsg2 robustly co-immunoprecipitated with WT-Pcad and I175D-Pcad, it was only weakly pulled down with W2A-Pcad (**Fig. 2a**). Quantification of the Dsg2/Pcad pull-down ratios revealed that I175D-Pcad retained ∼80% Dsg2 binding capacity of WT-Pcad, whereas W2A-Pcad retained only ∼25% Dsg2 binding capacity of WT-Pcad (**Fig. 2b**). This substantial reduction in WT-Dsg2:W2A-Pcad interactions in cells, further demonstrates the critical role of W2 in Pcad and Dsg2 association. Taken together, our AFM and co-IP data demonstrate that Pcad forms a Ca²⁺-dependent heterodimer with Dsg2 via a strand-swap trans interface mediated by Trp2.

**Figure 2.**
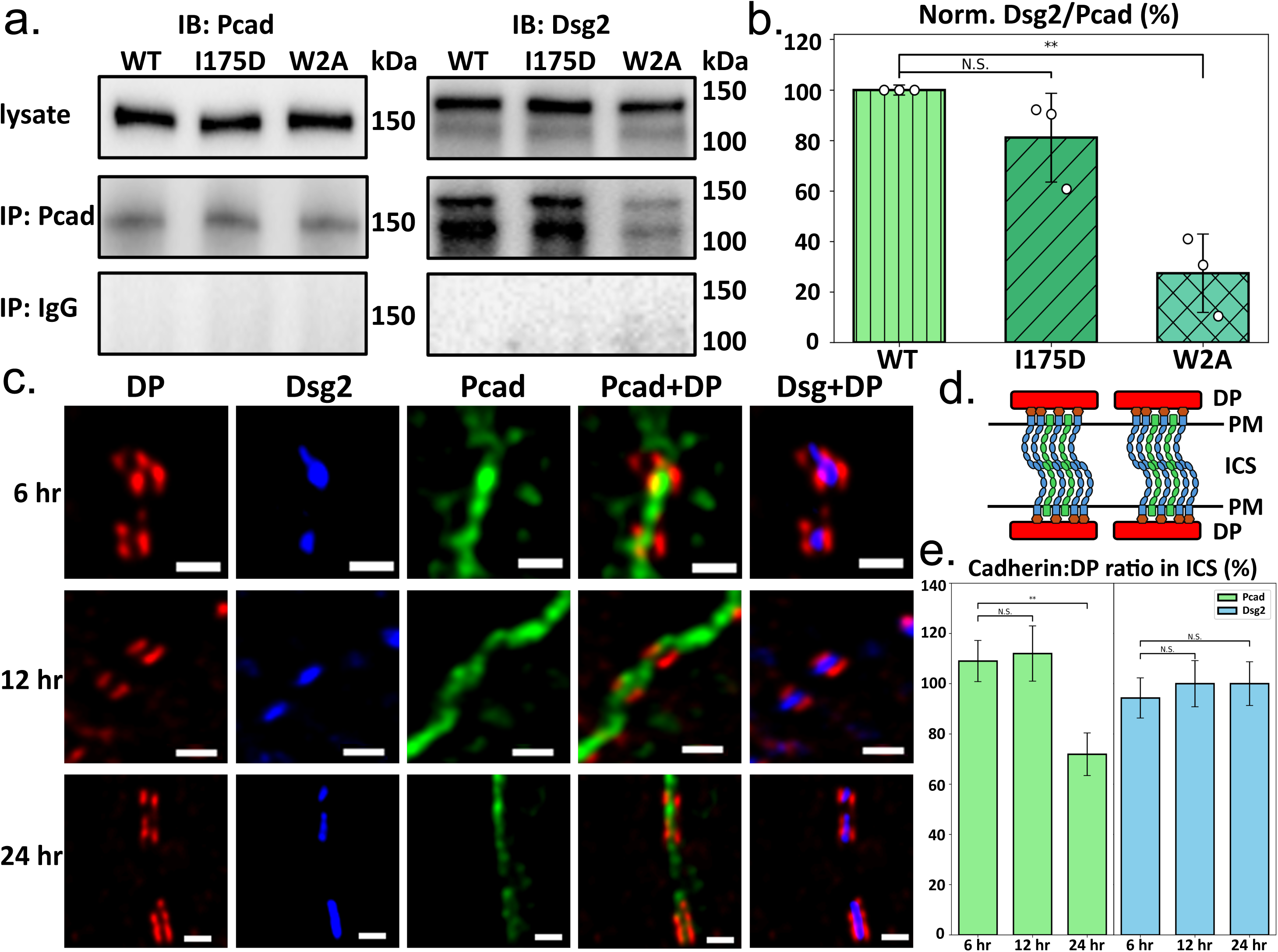
P-cadherin interacts with Desmoglien-2 to facilitate desmosome assembly. **(a)** Co-immunoprecipitation (co-IP) experiments show that the amount of WT-Dsg2 associated with W2A-Pcad is significantly reduced. Representative co-IP results are shown. Equal amounts of cell lysates were incubated with agarose beads, and no Pcad or Dsg2 was detected when IgG was used as a negative control for pull-down. **(b)** Quantification of the co-IP experiment. Bar plot depicts the Dsg2/Pcad ratio measured across three biological replicates, normalized to the values obtained for WT-Pcad. The statistical analysis was performed in comparison with data in WT condition. For I175D condition, the P-value is 0.137. For W2A conditions, the P-value is 1.26E-3. **(c)** Representative STED microscopy images of desmosomal regions in A431 cells following a calcium switch at 6, 12, or 24 hours. Scale bar: 500 nm. **(d)** Schematic representation of the characteristic “railroad track” structure of desmosomes. STED imaging resolves plaque-to-plaque spacing. Cadherin intensity (Pcad or Dsg2) within the railroad track region was quantified. **(e)** Relative intensity analysis normalized to average Dsg2 expression at 24 hours post-Ca²⁺ switch reveals that Pcad is highly expressed in early-stage (immature) desmosomes and decreases upon maturation. In contrast, Dsg2 expression remains relatively stable over time. Measurements from three independent biological replicates. For 6, 12, and 24 hour samples, n = 33, 30, and 35. Statistical analysis was performed in comparison with data in 6 hours samples. For Pcad relative expression: difference between 6 and 12 hours samples is not significant with a P-value of 0.834; difference between 6 and 24 hours samples is significant with a P-value of 2.55E-3. For Dsg2 relative expression: difference between 6 and 12 hours samples is not significant with a P-value of 0.784; difference between 6 and 24 hours samples is also not significant with a P-value of 0.766.

### Pcads are required for desmosome formation

To investigate the role of Pcad in desmosome assembly, we used stimulated emission depletion (STED) microscopy to examine Pcad’s localization within desmosomal regions at different stages of cell–cell junction formation in Ecad/Pcad KO cells rescued with either WT-Pcad, W2A-Pcad, or I175D-Pcad. We grew the cells to confluency and then used a calcium-switch to reset desmosome assembly. At defined time points following the calcium switch (6 hours, 12 hours, and 24 hours), cells were fixed and immunostained with antibodies against mCherry (for Pcad), Dsg2, and desmoplakin (DP) to assess expression and colocalization within desmosomal regions. While we also attempted the imaging experiments at an earlier timepoint (3 h), desmosomes at this early timepoint could not be unambiguously identified (Supplementary Fig. 1a).

Because DP is a core component of desmosomes and forms a characteristic discontinuous “railroad track” pattern ^19–21^, we used this morphological feature to identify desmosomes and quantify Pcad and Dsg2 localization (**Fig. 2c**, **2d**). Since prior studies have shown that Dsg2 expression remains relatively stable throughout the desmosome assembly process ^8,22^, we used Dsg2 intensity at the 24 hour time point (mature desmosomes) as a reference for relative quantification. Consistent with previous findings, Dsg2 levels remained stable (∼95% relative intensity) throughout the 24-hour observation period (**Fig. 2e**). In contrast, Pcad signal was initially higher than that of Dsg2 at 6 hours and continued to increase until 12 hours, prior to desmosome maturation. By 24 hours, Pcad levels had decreased to approximately 60% of their peak intensity (**Fig. 2e**). In contrast, previous studies have shown that while Ecad expression peaked at the onset of desmosome formation, it was rapidly downregulated within 3 hours ^8^. These data suggest that Pcad is enriched at nascent desmosomes and may play a more robust role in desmosome formation than Ecad.

To further investigate the role of Pcad in desmosome initiation, we tested whether its presence is required for Dsg2 and DP recruitment to cell–cell junctions. Confluent monolayers of A431 Ecad/Pcad KO cells rescued with either WT-Pcad, I175D-Pcad, or W2A-Pcad were exposed to a calcium switch and fixed 3 hours, 6 hours, and 24 hours post-Ca²⁺ reintroduction. The cells were stained for mCherry (i.e. Pcad), Dsg2, and DP (**Fig. 3a**). We first assessed colocalization between Pcad and Dsg2/DP. While colocalization of I175D-Pcad with both Dsg2 and DP was similar to WT-Pcad, W2A-Pcad exhibited reduced colocalization with both Dsg2 and DP compared to WT or I175D Pcad (**Figs. 3b, 3c**). These results corroborate our earlier co-IP data, confirming that the W2A mutation impairs Pcad’s association with desmosomal components. Next, we evaluated membrane recruitment of Dsg2 and DP by using Pcad localization at the cell border as a proxy for membrane boundaries. We calculated the membrane-associated fraction of Dsg2 and DP by quantifying the proportion of membrane-localized signal relative to total cellular signal (**Fig. 3a**). In cells expressing WT-Pcad or I175D-Pcad, Dsg2 and DP exhibited progressive enrichment at the membrane over time, consistent with successful recruitment during desmosome assembly (**Fig. 3e**, **3f**). In contrast, W2A-Pcad expressing cells showed impaired recruitment of both Dsg2 and DP (**Fig. 3e**, **3f**). Taken together, these findings demonstrate that Pcad is dynamically enriched at desmosomal junctions and facilitates the early recruitment of desmosomal components such as Dsg2 and DP. This recruitment critically depends on the strand-swap trans interface of Pcad and is impaired in the W2A mutant.

**Figure 3.**
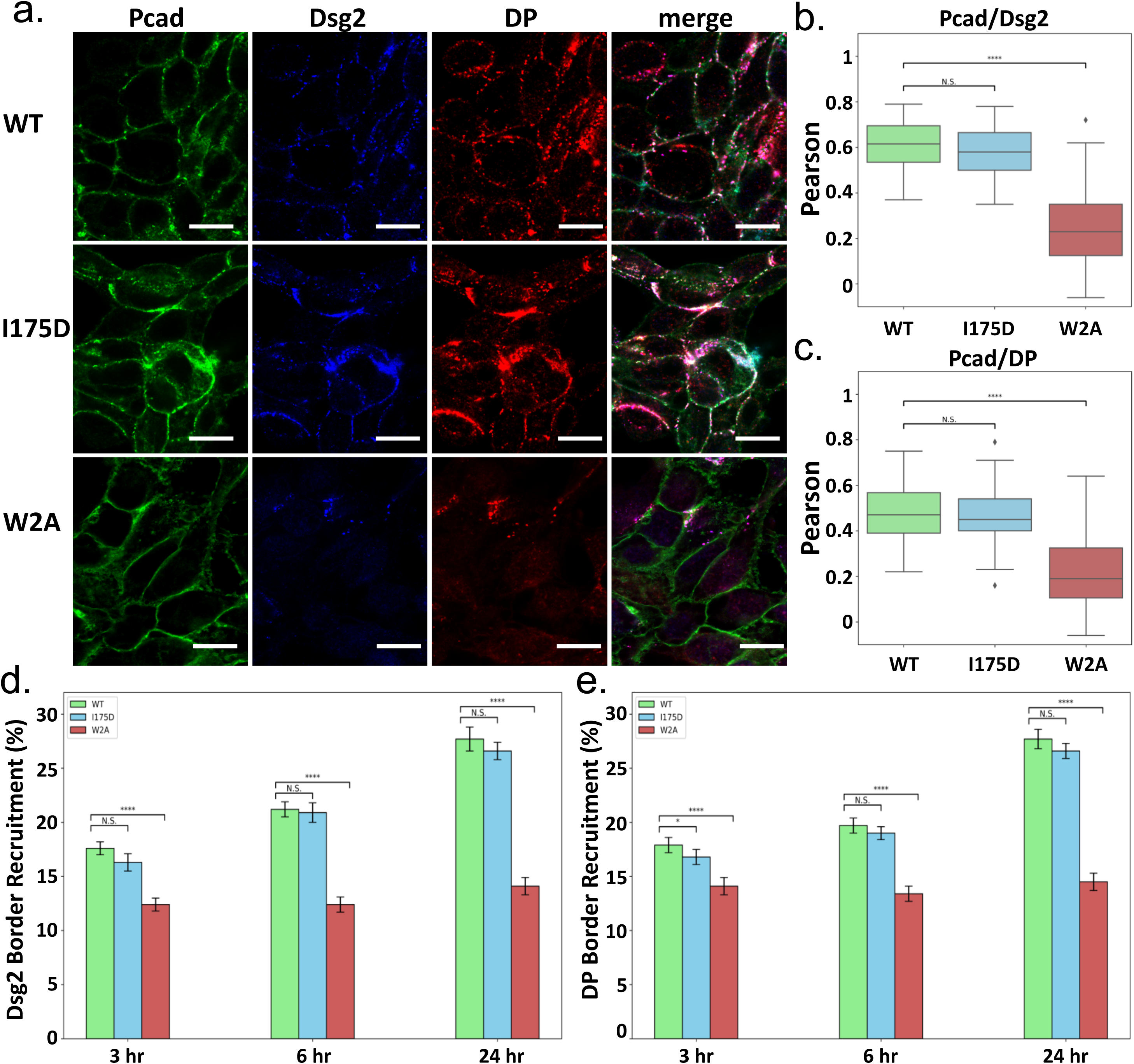
P-cadherin:Desmoglien-2 strand-swap interaction promotes Desmoglien-2/Desmoplakin recruitment to cell borders. **(a)** Immunofluorescence analysis of desmosomal component recruitment to cell borders in A431 cells fixed at 24 hours after a calcium-switch. Pcad (green), Dsg2 (blue), DP (red). Reduced Dsg2 and DP localization to cell borders is observed in cells expressing W2A-Pcad. Cells expressing I175D-Pcad show no significant difference in Dsg2/DP recruitment compared to cells expressing WT-Pcad. Scale bar: 10 µm. **(b–c)** Colocalization between Pcad and desmosomal components—Dsg2 or DP—is preserved in cells expressing either WT-Pcad or I175D-Pcad but is disrupted in cells expressing W2A-Pcad. For WT-Pcad, I175D-Pcad, and W2A-Pcad conditions, the numbers of cells were 54, 51, and 55 respectively, from three independent biological replicates.. Statistical analysis was performed in comparison with data in WT condition. For I175D condition, the P-value is 0.127 for Dsg2 and 0.488 for DP. For W2A conditions, the P-value is 2.92E-24 for Dsg2 and 3.25E-15 for DP. **(d–e)** Quantification of Dsg2 and DP recruitment to cell borders demonstrates that expression of W2A-Pcad (red) significantly impairs desmosomal component localization to cell border compared to WT-Pcad (green) or I175D-Pcad (blue). Measurements from three independent biological replicates. For the WT condition, the n for 3 hours, 6 hours, and 24 hours samples were 52, 60, and 55, respectively. For the I175D condition, the n for 3 hours, 6 hours, and 24 hours samples were 53, 55, and 48, respectively. For the W2A condition, the n for 3 hours, 6 hours, and 24 hours samples were 52, 48, and 53, respectively. Statistical analyses were performed in comparison with WT condition at each time point. For the Dsg2 recruitment level at 3 hours, the P value between WT-Pcad and I175D-Pcad is 0.786 and the P-value between WT-Pcad and W2A-Pcad is 9.14E-11. For the Dsg2 recruitment level at 6 hours, the P value between WT-Pcad and I175D-Pcad is 0.687 and the P-value between WT-Pcad and W2A-Pcad is 3.76E-14. For the Dsg2 recruitment level at 24 hours, the P value between WT-Pcad and I175D-Pcad is 0.744 and the P-value between WT-Pcad and W2A-Pcad is 8.57E-16. For the DP recruitment level at 3 hours, the P value between WT-Pcad and I175D-Pcad is 0.151 and the P-value between WT-Pcad and W2A-Pcad is 2.29E-07. For the DP recruitment level at 6 hours, the P value between WT-Pcad and I175D-Pcad is 0.453 and the P-value between WT-Pcad and W2A-Pcad is 5.36E-09. For the Dsg2 recruitment level at 24 hours, the P value between WT-Pcad and I175D-Pcad is 0.109 and the P-value between WT-Pcad and W2A-Pcad is 8.65E-19.

### Novel molecular interactions stabilize Pcad:Dsg2 S-dimers

Although the crystal structures of homophilic S-dimers formed by Pcad ^23^, Ecad ^15^, and Dsg2 ^17^ have been resolved previously, the structure of a strand-swap heterodimer between Pcad and Dsg2 remains unknown. To investigate the molecular interactions underlying heterophilic binding, we computationally modeled a Pcad:Dsg2 S-dimer by aligning the EC1-2 domains of Pcad against one protomer of the Dsg2 homodimer and replacing it (**Fig. 4a**). For comparative analysis, we also aligned the EC1-2 domains of Ecad to the same protomer of Dsg2 homodimer to assess the potential barrier for Ecad:Dsg2 S-dimer formation (**Fig. 4b**). To minimize steric clashes at the dimer interface, the alignment was primarily guided by the EC1 domains of Pcad/Ecad and Dsg2. We then performed molecular dynamics (MD) simulations on both the Pcad:Dsg2 and Ecad:Dsg2 aligned S-dimers, using previously established protocols ^24,25^. Each system was simulated in five independent replicates for 50 ns until equilibrium was reached (**Supplementary Fig. 3**). Notably, during the MD simulations, the Root Mean Square Deviation (RMSD) relative to the initial structures at the start of the simulation was much higher in all Ecad:Dsg2 sets compared to Pcad:Dsg2 sets (**Supplementary Fig. 3**), suggesting that S-dimers formed by Ecad:Dsg2 were less stable than those formed by Pcad:Dsg2 (**Supplementary Fig. 4**). To assess the stability of the strand-swap interface, we measured the center of mass (COM) distance between the Trp2 residue of Pcad or Ecad and the Dsg2 hydrophobic pocket composed of Ile24, Tyr38, Ala82, and Leu94 ^17^. The Trp2 from Pcad remained stably buried in the Dsg2 pocket throughout the simulation, whereas Trp2 from Ecad frequently detached from the pocket (**Fig. 4c**).

**Figure 4.**
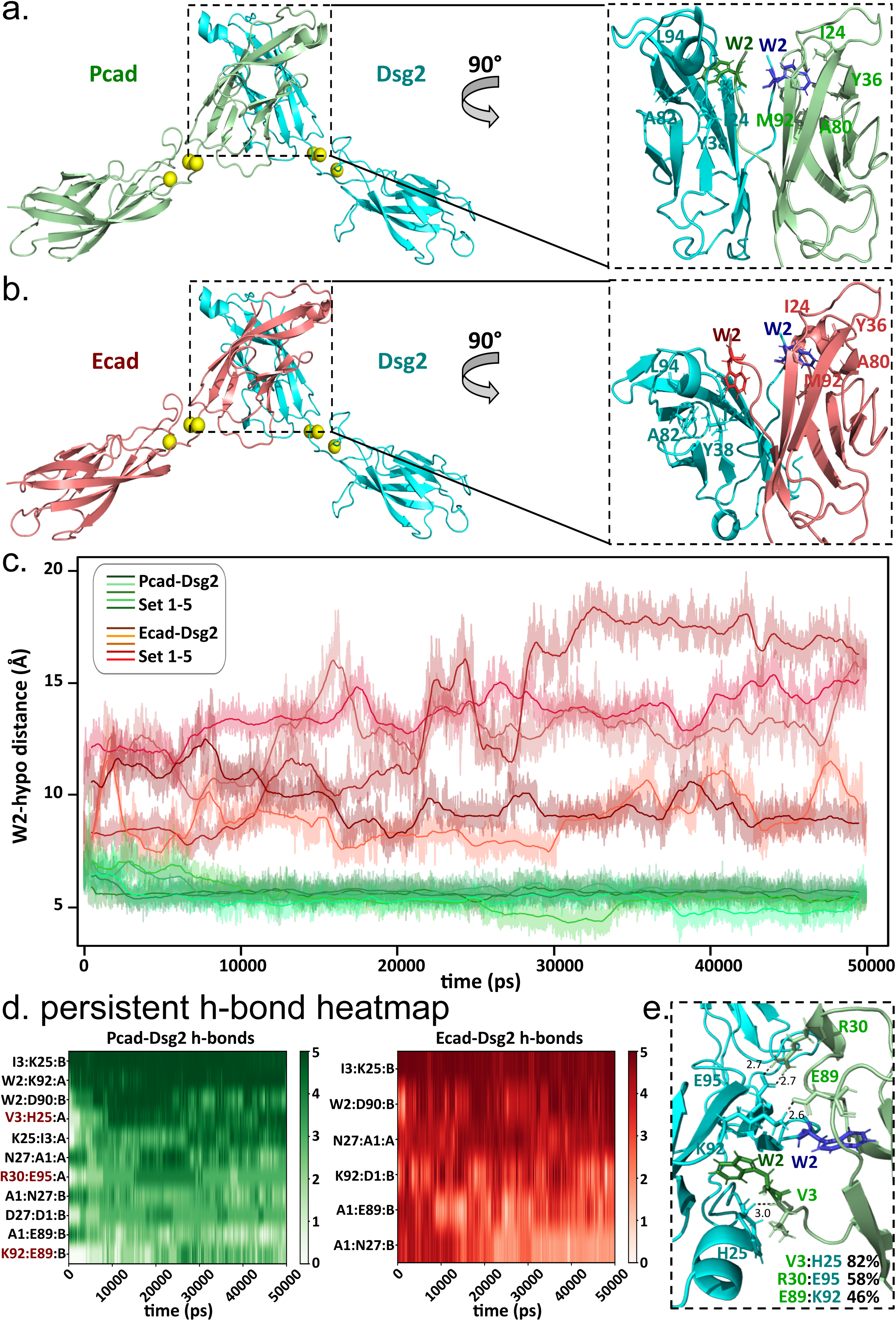
P-cadherin and Desmoglien-2 form stable strand-swap dimers. **(a)** Simulation setup for the aligned structure of the Pcad:Dsg2 strand-swap dimer. Right panel: zoomed-in view of a representative final frame from the Pcad:Dsg2 simulation shows stabilization of opposing Trp2s inserted into their complementary hydrophobic pocket. **(b)** Simulation setup for the aligned structure of the Ecad:Dsg2 strand-swap dimer. Right panel: zoomed-in view of a representative final frame from the Ecad:Dsg2 simulation shows destabilization of Ecad’s Trp2 and its dissociation from Dsg2’s complementary hydrophobic pocket. **(c)** Comparison of the paired distances between the centers of mass (COMs) of Trp2 and the complementary hydrophobic pocket across five MD simulation replicas reveals consistent stabilization of Pcad’s Trp2, whereas Ecad’s Trp2 consistently dissociates from the hydrophobic pocket. **(d)** Frequency heat map of persistent electrostatic interactions observed across five MD simulations. The Pcad:Dsg2 (green map) S-dimer exhibits 11 persistent electrostatic interactions, whereas the Ecad:Dsg2 (red map) dimer shows only six persistent electrostatic interactions. Three novel interactions—V3:H25, R30:E95, and E89:K92— unique to the Pcad:Dsg2 simulations are highlighted. **(e)** Zoomed-in view of the strand-swapping trans interface showing the three novel electrostatic interactions. V3:H25 (present in 82% of total frames) stabilizes the Pcad β-strand. R30:E95 (58%) and E89:K92 (46%) contribute to the stabilization of the outer interface of the Pcad:Dsg2 S-dimer.

To further evaluate interface stability, we analyzed electrostatic interactions across all five simulation replicates **(Supplementary Fig. 5 and Supplementary Fig. 6**). Interactions persisting for more than 30% of frames in a given trajectory were classified as persistent, and we compiled these interactions across all 5 replicates into a heatmap (**Fig. 4d**). Notably, the Pcad:Dsg2 complex exhibited a higher number of persistent molecular interactions than the Ecad:Dsg2 complex, suggesting that Pcad is more structurally compatible with Dsg2 for strand-swap dimerization. Three unique molecular interactions were identified as consistently persistent only in the Pcad:Dsg2 dimer: Val3:His25, Arg30:Glu95, and Glu89:Lys92 (**Fig. 4e**), each persisting in over 40% of total simulation frames. The Val3:His25 hydrogen bond, located adjacent to Trp2, appears to stabilize Trp2 within the Dsg2 pocket. Meanwhile, the Arg30:Glu95 and Glu89:Lys92 interactions, although located on the periphery of the trans interface, likely contribute to overall complex stability. Interestingly, although Val3, Arg30, and Glu89 are conserved in Ecad, it fails to form the same electrostatic interactions with Dsg2 because the surrounding residues in Ecad sterically hinder formation of these hydrogen bonds and salt bridges. In summary, our MD simulations reveal that Pcad forms a more stable strand-swap heterodimer with Dsg2 than Ecad, supported by persistent Trp2 insertion and enriched intermolecular interactions.

### β-strand hinge is a key determinant in Pcad:Dsg2 S-dimer formation

The anchor residue Trp2 which is swapped in an S-dimer, resides on an N-terminal β-strand ^15^. Structurally, this swapped β-strand can be subdivided into three regions based on the presence of a conserved proline hinge (Pro5-Pro6 in Ecad and Ala5-Pro6 in Pcad) ^26^: residues 1–4 comprise the pre-hinge strand (PH), residues 5–6 form the hinge region, and residues 7–10 constitute the after-hinge strand (AH) (**Fig. 5a**). Prior studies demonstrated that mutating both Pro5-Pro6 in mouse Ecad to Ala results in an extended β-strand interface with enhanced hydrogen bonding to adjacent strands, leading to increased binding affinity ^26^. In contrast, the rigid proline hinge in WT Ecad restricts β-strand extension and limits affinity. A sequence alignment of human Dsg2 and human classical cadherins shows that only Pcad and Dsg2 share an Ala5–Pro6 (AP) hinge motif (**Fig. 5b**). We therefore hypothesized that the AP hinge would confer increased flexibility, potentially promoting formation of a more favorable S-dimer between Pcad and Dsg2. To test this hypothesis, we computationally mutated the Ala5 in Pcad to Pro (A5P-Pcad mutant) within the aligned Pcad:Dsg2 heterodimer structure and performed MD simulations to assess structural and interaction changes. As in previous experiments, five independent replicates were simulated.

**Figure 5.**
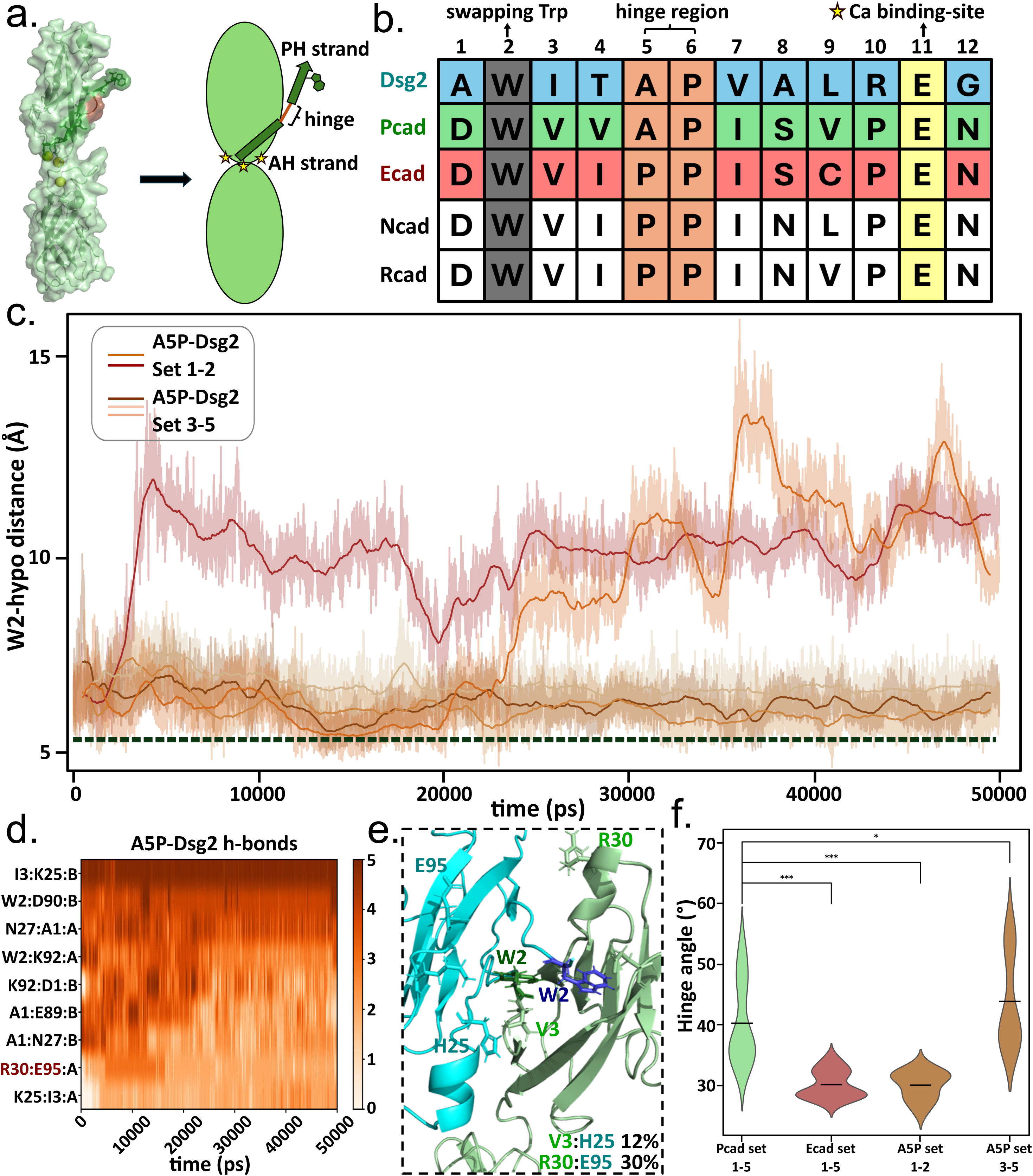
Hinge region is critical for P-cadherin:Desmoglein-2 strand-swap dimerization. **(a)** Structural representation of the Pcad EC1–2 domains highlighting the Ala5-Pro6 hinge region (brown), which divides the β-strand into the pre-hinge (PH), hinge, and after-hinge (AH) strands. **(b)** Sequence alignment analysis of human classical cadherins (Ecad, Pcad, Ncad, and Rcad) and human Dsg2 indicates that both Pcad and Dsg2 share a conserved Ala5-Pro6 hinge region, whereas all other classical cadherins possess a conserved Pro5-Pro6 hinge region. **(c)** Comparison of the distances between the COMs of Pcad-Trp2 and the complementary Dsg2-hydrophobic pocket across five MD simulation replicas of the A5P-Pcad:WT-Dsg2 complex. Trp2 in the A5P-Pcad mutant is destabilized in two replicas. Although Trp2 remains inserted in the remaining three replicas, the stabilized distances are greater than the corresponding distances in WT-Pcad:WT-Dsg2. The dashed black line represents the average Trp2-to-pocket distance in WT-Pcad. **(d)** Frequency heat map of persistent electrostatic interactions observed in the A5P-Pcad:WT-Dsg2 simulations shows that only the R30:E95 interaction remains consistently present. **(e)** Zoomed-in view of the destabilized strand-swapping trans interface in the A5P-Pcad:WT-Dsg2 complex. V3:H25 is present in only 12% of total frames, R30:E95 in 30% of total frames, and E89:K92 in less than 1% of total frames. **(f)** Hinge angle analysis reveals that the two destabilized A5P-Pcad:WT-Dsg2 simulations and the five WT-Ecad:WT-Dsg2 simulations exhibit similar hinge angles and reduced flexibility in the hinge region. In contrast, the three stabilized A5P-Pcad:WT-Dsg2 simulations resemble the five WT-Pcad:WT-Dsg2 simulations, both showing similar hinge angles and greater hinge flexibility. Statistical analyses were performed in comparison with the five WT-Pcad:WT-Dsg2 simulations. For the five WT-Ecad:WT-Dsg2 simulations, the P-value is 1.44E-15. For the two destabilized A5P-Pcad:WT-Dsg2 simulations, the P-value is 5.73E-14. For the three stabilized A5P-Pcad:WT-Dsg2 simulations, the P-value is 3.76E-02.

To evaluate interface stability, we measured the center of mass (COM) distance between Trp2 of A5P-Pcad and the Dsg2 hydrophobic pocket. In three of five simulations, Trp2 remained stably docked, while in two simulations, it disengaged from the pocket (**Fig. 5c**). Even in simulations where Trp2 remained bound, the average COM distance (∼7.2 Å) was notably larger than in WT-Pcad:Dsg2 simulations (∼5.5 Å), indicating reduced stability (**Fig. 5c**). To uncover the molecular mechanism underlying this destabilization, we analyzed persistent electrostatic interactions across the simulations (**Supplementary Fig.7**) and generated a persistent interaction heatmap (**Fig. 5d**). Our analysis revealed that the A5P mutation disrupted key interactions previously observed in the WT-Pcad:Dsg2 complex: the occurrence of the Val3:His25 hydrogen bond dropped to 12% of frames, the Glu89:Lys92 molecular interaction appeared in <1% of frames, and the Arg30:Glu95 interaction was reduced to ∼30% of the frames (**Fig. 5e**). These disruptions underscore the importance of hinge flexibility in maintaining favorable electrostatic contacts at the strand-swap interface.

Since prior studies have suggested that altering hinge residues can also affect the flexibility of the β strand ^26^, we next measured the angle between the principal axes of the PH and AH strands across the final 10 ns of each simulation. The hinge angles remained consistently small in all five Ecad:Dsg2 simulations and in the two A5P-Pcad:Dsg2 simulations where Trp2 was displaced (**Fig. 5f**). However, the three A5P-Pcad:Dsg2 simulations with retained Trp2 as well as all five WT-Pcad:Dsg2 simulations showed a much larger hinge angle (**Fig. 5f**), supporting the notion that hinge geometry influences interface stability.

To experimentally validate these findings, we generated complete ectodomain mutants of A5P-Pcad for single-molecule AFM binding assays (**Fig. 6a**). As expected, the mutation disrupted binding between A5P-Pcad and WT-Dsg2 (4.5% in Ca²⁺ and 3.1% in EGTA; **Fig. 6b**). Furthermore, interactions between A5P-Pcad and W2A-Dsg2 was reduced to nonspecific levels (2.0% in Ca²⁺ and 2.2% in EGTA; **Fig. 6b**). In summary, our data showed that the A5P mutation altered the β-strand hinge geometry in Pcad, disrupting critical electrostatic interactions with Dsg2 and impairing S-dimer formation both in silico and in vitro.

**Figure 6.**
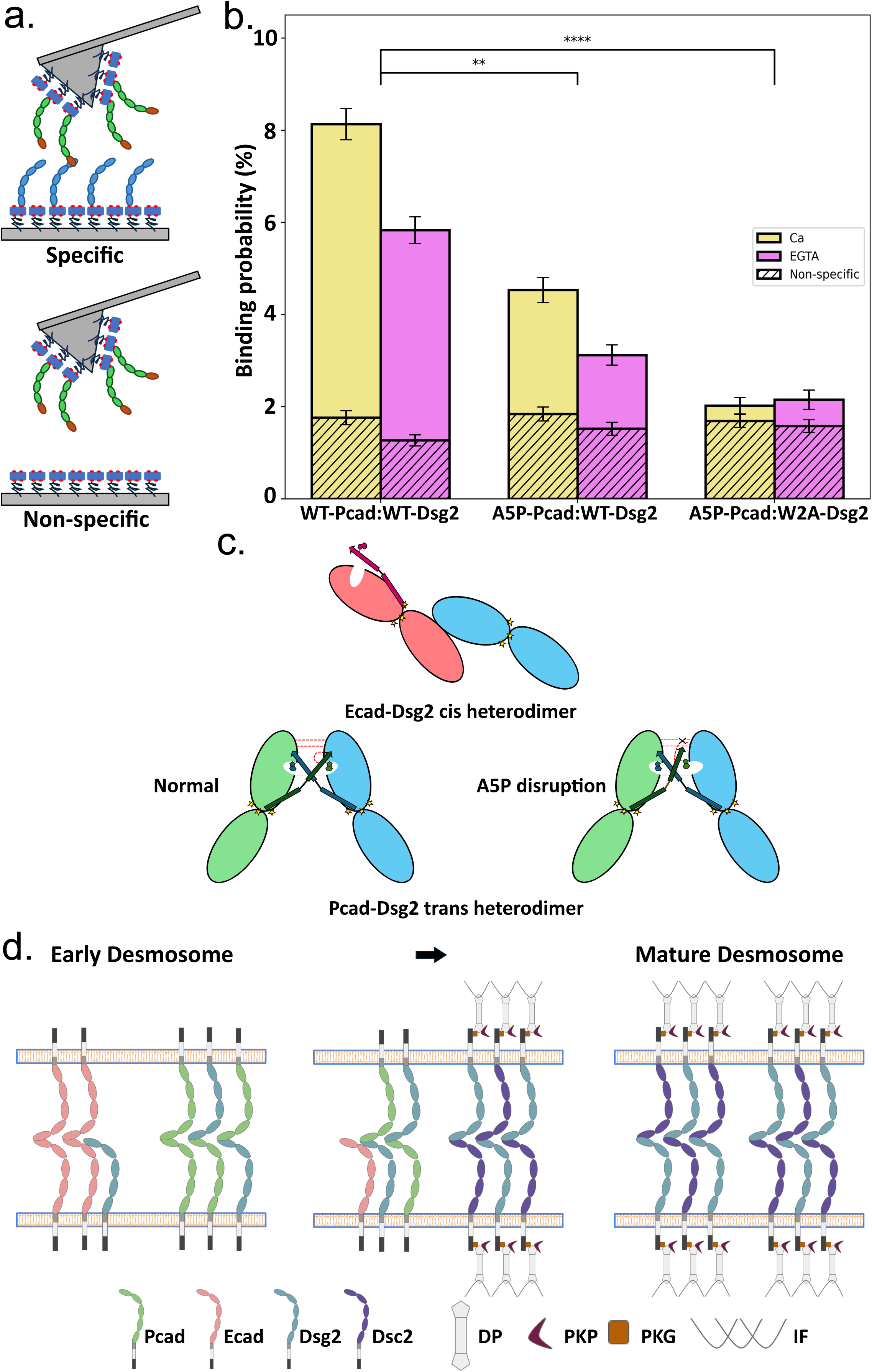
A5P mutation disrupts P-cadherin:Desmoglein-2 heterophilic interaction in vitro. **(a)** Schematic representation of the single-molecule AFM binding assay used to evaluate A5P-Pcad:Dsg2 heterophilic interactions. **(b)** The binding probability between A5P-Pcad and WT-Dsg2 is significantly lower than that observed between WT-Pcad and WT-Dsg2. The interaction between A5P-Pcad and W2A-Dsg2 indicates that the binding probability is reduced to the level of non-specific interactions. For all specific measurements, n = 6075, obtained from three independent replicates. For all non-specific measurements, n = 4050, obtained from four independent replicates. The statistical analyses were all performed in comparison with WT-Pcad:WT-Dsg2 group. For A5P-Pcad:WT-Dsg2, the P-value is 1.08E-3. For A5P-Pcad:W2A-Dsg2, the P-value is 1.30E-5. Error bars represent the standard error calculated using bootstrapping with replacement. **(c)** Schematic illustration of the molecular mechanisms underlying WT-Ecad:WT-Dsg2 interactions (cis dimers), WT-Pcad:WT-Dsg2 interactions (trans S-dimers), and the A5P mutation–mediated disruption of Pcad:Dsg2 binding. **(d)** Schematic representation of the roles of Pcad and Ecad in desmosome assembly. Ecad forms transient cis heterodimers with Dsg2, whereas Pcad forms more persistent trans heterodimers with Dsg2. As the desmosome matures, both Ecad:Dsg2 heterodimers and Pcad:Dsg2 heterodimers dissociate, allowing Dsg2 and Dsc2 to interact and form a stable adhesive complex.

## Discussion

Using a combination of in vitro single-molecule experiments, cell-based functional assays, and atomistic simulations, we demonstrate that Pcad interacts with Dsg2 in a robust, Ca²⁺-dependent S-dimer conformation and promotes desmosome assembly. Our findings extend existing models for desmosome assembly by accounting for mechanistic differences between Ecad and Pcad (**Fig. 6d**). Unlike Ecad, which transiently binds to Dsg2 in a cis orientation during early desmosome formation ^8^ (**Fig. 6c**), we show that Pcad and Dsg2 form more stable trans strand-swap interactions which serve as a structurally robust mechanism for recruiting desmosomal components to nascent junctions (**Fig. 6c**, **6d**). As the desmosome matures, Pcad is excluded from the final desmosomal core, consistent with our observation that Pcad signal declines at later stages of junction maturation; albeit much more slowly compared to Ecad ^8^. Thus, our data suggests that Ecad and Pcad facilitate desmosome formation through mechanistically distinct but functionally convergent pathways. Our data highlights the molecular diversity of cadherin-mediated adhesion. More broadly, our findings support a model in which the heterodimerization of classical and desmosomal cadherins is regulated in space and time by interface geometry, local flexibility, and interaction energetics—principles that are likely to apply to other dynamic adhesive systems in development, wound healing, and cancer progression.

Our data suggests that the Ala5-Pro6 hinge region in the β strands of Dsg2 and Pcad are a critical determinant of heterophilic S-dimer formation. Prior work has shown that the β strand hinge in Ecad, which is made up of Pro5-Pro6 residues, plays a crucial role in strand-swap homodimer formation, and that mutating one or both Pro residues can enhance Ecad–Ecad binding affinity ^26^. These studies suggest that alanine substitutions increase hinge flexibility and promote interactions with the opposing β-strand ^26^. Among all Dsg isoforms, Dsg2 uniquely contains an Ala5-Pro6 hinge, while the other isoforms contain an Phe5-Ala6 hinge (**Supplementary Fig. 8a**). Similarly, all isoforms of Dsc contain an Ile5-Pro6 hinge which likely has limited flexibility due to Ile’s bulky hydrophobic side chain (**Supplementary Fig. 8a**). Similarly, of the four well-characterized human classical cadherins—Ecad, Pcad, Ncad, and Rcad—, only Pcad has an Ala5-Pro6 hinge (**Fig. 5b**). Consequently, unlike Ecad which cannot interact with Dsg2 in a trans conformation ^8^, Pcad and Dsg2 form trans S-dimers. More broadly, we anticipate that the principal of hinge flexibility can be used to predict heterophilic strands-swapping between other desmosomal and classical cadherins.

Unlike human Pcad, its murine ortholog does not contain an Ala5-Pro6 hinge motif but instead has a Pro5-Pro6 hinge (**Supplementary Fig. 8b**). In contrast, the Ala5-Pro6 hinge is conserved in primate Pcads from green monkey, gorilla, and chimpanzee, indicating evolutionary conservation among primates (**Supplementary Fig. 8b**). The absence of the Ala5-Pro6 hinge motif in mouse Pcad may be a limitation when murine models are used to study demosome assembly.

Canine Ecad, which shares high structural homology with Pcad, forms a Ca²⁺-independent heterophilic cis dimer with Dsg2 via a conserved Leu175 residue ^8^. In contrast, Pcad and Dsg2 do not form cis dimers but instead form Ca²⁺-dependent trans S-dimers. The different heterophilic cis dimerization capabilities of Pcad and Ecad may stem from the different electrostatic properties of their binding interfaces. While Pcad exhibits a localized positively charged surface (**Supplementary Fig. 9b**), Ecad displays a negatively charged patch (**Supplementary Fig. 9c**) on its cis binding interface. The opposing Dsg2 surface, identified through cis-interface structural alignment, is also positively charged ^27,28^ (**Supplementary Fig. 9a**). The electrostatic mismatch between Pcad and Dsg2 may explain why Pcad is unable to engage Dsg2 via a cis orientation and instead favors trans-mediated interactions.

Previous studies reported Pcad and Dsg2 co-localization at intercellular contacts in C32 melanoma cells ^29^. Our co-immunoprecipitation experiments validated interaction between Pcad and Dsg2 during desmosome assembly while super-resolution imaging, visualized Pcad localization within nascent desmosomes. Pcad’s integration into nascent desmosomes may imply a role in promoting junctional recruitment before the full maturation of the desmosomal complex. Given that Ecad’s cis interaction with Dsg2 is known to be less robust than trans interactions ^8,30^, the sustained presence of Pcad during desmosome assembly suggests that it plays a more specific stabilizing role in supporting Dsg2 function. Temporal quantification confirms that the Pcad:Dsg2 interaction is more persistent during early stages of desmosome assembly compared to Ecad:Dsg2. The formation of robust Pcad:Dsg2 S-dimers may also explain previous studies showing that Pcad substitutes for Dsg3 during the desmosome assembly process ^2,10^.

Our confocal immunofluorescence microscopy analysis of Dsg2 and DP border recruitment further confirms the role of Pcad in desmosome assembly and demonstrates that S-dimer–deficient Pcads abolish Dsg2 and DP recruitment to cell borders during desmosome formation (**Fig. 3**). While a recent study showed that W2A-Pcad is turned over more rapidly than WT-Pcad ^31^, this is unlikely to influence desmosomal component recruitment since the same study also confirmed that the membrane localization of WT-Pcad and W2A-Pcad were similar. Indeed, our western blots demonstrate that WT-Pcad, I175D-Pcad, and W2A-Pcad are expressed at similar levels (**Supplementary Fig. 2a**). Furthermore, Dsg2 expression also remains comparable across WT-Pcad, I175D-Pcad, and W2A-Pcad cells (**Fig. 2a**). This suggests that comparable amounts of WT or mutant Pcad are present on the cell surface to bind Dsg2 and that W2A-Pcad decreases Dsg2 and DP recruitment since it is unable to form heterophilic S-dimers with Dsg2.

Since the cytoplasmic proteins plakoglobin and β-catenin are close homologs with a shared ability to bind the cytoplasmic tails of cadherins ^32,33^, it has previously been suggested that the Ecad–plakoglobin complex in A431 cells facilitates desmosome assembly at nascent contacts ^34^. Because Pcad, like Ecad, recruits β-catenin during adherens junction assembly ^4^, it is reasonable to hypothesize that Pcad can also facilitate desmosome assembly by directly engaging plakoglobin at early contacts, thereby positioning plakoglobin for association with desmosomal cadherins. In this scenario, potential heterophilic Pcad:Dsg2 interactions at the membrane would provide an extracellular cue that promotes plakoglobin recruitment and desmosome maturation.

Using single molecule AFM measurement we demonstrated that W2A-Pcad:W2A-Dsg2 did not interact; their binding probability was similar to non-specific binding. In contrast, the binding probabilities for W2A-Pcad:WT-Dsg2 and for WT-Pcad:W2A-Dsg2 were roughly 50% of the WT-Pcad:WT-Dsg2 binding probability. This suggests that W2A-Pcad:WT-Dsg2 and WT-Pcad:W2A-Dsg2 form asymmetric strand-swap dimers, where the WT-cadherin inserts its Trp2 residue into the binding pocket on its W2A-cadherin binding partner without reciprocal strand-swapping. A similar asymmetric strand swapping has been recently described for the binding of WT-Ecad and W2A-Ecad ^35^.

While prior crystallographic studies have resolved strand-swapped homodimers of Pcad^23^, Ecad^15^, and Dsg2^17^, a crystal structure for the Pcad:Dsg2 S-dimer does not exist. Consequently, we generated the structures of Pcad:Dsg2 and Ecad:Dsg2 S-dimers based on structure alignment followed by MD simulations. While our analysis revealed that the Pcad:Dsg2 heterodimer possesses a distinct set of stabilizing interactions, it is important to recognize that the predicted interactions are highly dependent on the accuracy of the structural models. Fortunately, the structures of both Pcad and Dsg2 ectodomains have been resolved using X-ray crystallography. More importantly, we experimentally confirmed the predictions of our atomistic MD simulations using single molecule AFM measurements.

The interplay of Dsg2 and Pcad is implicated in several cancers. Dsg2 expression is known to be upregulated in lung ^36,37^ and skin cancers ^38^ and downregulated in gastric cancer ^39^, often correlating with impaired cell adhesion and desmosome remodeling ^40,41^. Additionally, Pcad has been shown to modulate desmosome-associated signaling pathways that influence malignant transformation ^42,43^. Consequently, the Pcad–Dsg2 S-dimers may serve as a therapeutic biomarker or intervention point in these disease states. Our findings that Pcad and Dsg2 form stable S-dimers which rely on the flexibility of the Ala5-Pro6 hinge may also have therapeutic implications since this hinge could serve as a target for small molecules or antibodies aimed at modulating adhesion. Finally, conformation specific antibodies that target desmosome assembly can be designed by targeting Pcad:Dsg2 S-dimers; similar antibodies such as CQY684^31^, 19A11^24^ and 66E8^25^, have been previously designed to target Pcad and Ecad adhesive conformations.

## Materials and Methods

### Generation and purification of WT, I175D, W2A and A5P Pcad Ectodomains, and WT and W2A Dsg2 Ectodomains

The extracellular domains of human Pcad (residues 1–654) and Dsg2 (residues 1–609) were engineered with C-terminal Avi- and 6×His-tags and cloned into the pcDNA3.1(+) vector, as previously described^24,25,31^. Site-directed mutagenesis (Q5 Site-Directed Mutagenesis Kit, New England Biolabs) was used to generate the I175D, W2A, and A5P Pcad mutants, as well as the W2A Dsg2 mutant. These constructs were transfected into Expi293 suspension cells using the ExpiFectamine Kit (Thermo Fisher). After a 5-day expression period, culture supernatants containing the secreted target proteins were harvested and treated with protease inhibitors (Thermo Fisher Scientific) to prevent degradation.

All proteins (wild-type and mutants) were purified using Ni-NTA agarose beads (Qiagen) via HPLC columns (Bio-Rad). The beads were first washed with deionized (DI) water, followed by a binding buffer (20 mM HEPES, 500 mM NaCl, 1 mM CaCl₂, pH 7.5) containing the conditioned media from the previous step. Proteins bound to the Ni-NTA resin were then washed with biotinylation buffer (25 mM HEPES, 5 mM NaCl, 1 mM CaCl₂, pH 7.5) and biotinylated using the BirA enzyme (BirA500 Kit, Avidity). Elution was performed with a buffer containing 500 mM NaCl, 1 mM CaCl₂, and 200 mM imidazole (pH 7.5). The eluents were dialyzed against storage buffer (10 mM Tris-HCl, 100 mM NaCl, 10 mM KCl, 2.5 mM CaCl₂, pH 7.5) to remove imidazole. For long-term storage, 20% glycerol was added to the samples, which were then flash-frozen in liquid nitrogen and stored at –80 °C.

### Single-molecule AFM experiments

Single-molecule AFM experiments were performed following established protocols ^24,25,31^. Prior to functionalization, cantilevers (Hydra-2R-50N, AppNano) and glass coverslips (CS) were cleaned by overnight immersion in Piranha solution (30% H₂O₂, 70% H₂SO₄), followed by three rinses with deionized (DI) water. Coverslips were subsequently treated with 1 M KOH, rinsed again, and both cantilevers and coverslips were washed with acetone. Surface silanization was carried out by incubating the cleaned surfaces in 2% 3-aminopropyltriethoxysilane (Millipore Sigma) in acetone for 30 minutes at room temperature. The silanized surfaces were then functionalized with a mixture of 10% biotin-PEG-succinimidyl valerate and 90% mPEG-succinimidyl valerate (MW 5000, Laysan) in a solution containing 100 mM NaHCO₃ and 600 mM K₂SO₄ for at least 4 hours, followed by rinsing with DI water.

Prior to the experiments, surfaces were blocked overnight with 1 mg/mL BSA, then incubated with 0.1 mg/mL streptavidin (Invitrogen) for 30 minutes at room temperature. Purified protein was immobilized onto both cantilevers and coverslips by incubating with a 200 nM solution of the protein for 90 minutes. To minimize non-specific interactions, the protein-coated cantilevers and coverslips were incubated with 0.2 ng/mL D-biotin for 30 minutes.

AFM experiments were carried out using an Agilent 5500 AFM system equipped with a closed-loop scanner. Measurements were performed in Tris buffer (pH 7.5) containing 10 mM Tris-HCl, 100 mM NaCl, 10 mM KCl, and 2.5 mM CaCl₂. Spring constants of the cantilevers were determined using the thermal fluctuation method. Specific molecular interactions were identified by analyzing PEG force-extension curves, first using a freely jointed chain (FJC) model, followed by refinement with a worm-like chain (WLC) model using least-squares fitting.

### Stable Pcad rescued cell line construction

Full-length WT, I175D, and W2A Pcad constructs, each fused to a C-terminal mCherry tag, were transfected into Ecad/Pcad KO A431 cells. Stable single-cell clones expressing each construct were isolated by limiting dilution and subsequently expanded. Clonal populations were further purified by fluorescence-activated cell sorting (FACS), and monoclonal lines were established by seeding individual cells into collagen-coated 96-well plates. Cells were cultured in high-glucose Dulbecco’s Modified Eagle Medium (DMEM; Gibco) supplemented with 10% fetal bovine serum (FBS) and 1% penicillin–streptomycin.

### Co-immunoprecipitation (co-IP)

A431 cells were cultured in T75 flasks (Thermo Fisher Scientific) and harvested upon reaching confluence. After 48 hours, the cells were washed three times with pre-warmed phosphate-buffered saline (PBS; Invitrogen) and collected by scraping into 0.7 mL of PBS. The harvested cells were then centrifuged at 3,000 rpm for 10 minutes at room temperature to obtain cell pellets. Pellets were lysed in M2 lysis buffer (50 mM Tris-HCl, pH 7.5; 150 mM NaCl; 1% SDS; 1% Triton X-100) supplemented with 1% protease inhibitor cocktail (Sigma-Aldrich) and 0.1% Benzonase (Millipore Sigma). Lysates were flash-frozen in liquid nitrogen, thawed in a 37 °C water bath, and incubated with gentle inversion for 30 minutes at 4 °C. To ensure complete lysis, samples were sonicated twice for 1 minute. Insoluble debris was removed by centrifugation at 13,000 rpm for 30 minutes at 4 °C. Total protein concentration was determined using the Bio-Rad DC protein assay kit with BSA as the standard. To maintain consistency, a final protein concentration of 2 mg/mL was used for subsequent experiments.

For immunoprecipitation, magnetic Protein G beads (Thermo Fisher Scientific) were incubated with anti-mCherry rabbit monoclonal antibody (#43590, Cell Signaling Technology) for 30 minutes at room temperature. The antibody-conjugated beads were then mixed with 500 μL of cell lysate and incubated with rotation for 30 minutes at room temperature. Beads were washed three times with wash buffer (Thermo Fisher Scientific), and bound proteins were eluted by incubating with elution buffer for 5 minutes at room temperature.

### Western blot analysis

Samples were boiled at 95 °C for 10 minutes in SDS-PAGE sample buffer (Bio-Rad; 90% Laemmli + 10% 2-mercaptoethanol). Proteins were resolved on 4–15% polyacrylamide Mini-PROTEAN TGX gels (Bio-Rad) at 200 V for 30 minutes, then transferred to PVDF membranes (Bio-Rad) at 200 mA for 60 minutes on ice.

Membranes were blocked in 5% blocking buffer (PBS with 0.1% Tween 20 and 5% blotting-grade blocker) for 1 hour, followed by three washes with PBST. Blots were incubated with primary antibodies (1:1000 dilution) for 1 hour, then with secondary antibodies (1:5000 dilution) for an additional hour. Signal detection was performed using WesternBright ECL HRP substrate (Advansta).

### Stimulated Emission Depletion (STED) super resolution imaging

A431 cells were seeded onto 22 mm No. 1.5 glass coverslips coated with Bovine Collagen I (R&D Systems), placed in six-well plates, and cultured in a 5% CO₂ incubator at 37 °C for 48 hours. To ensure consistent desmosome maturation across all samples, a calcium switch protocol was applied. Cells were first rinsed with pre-warmed phosphate-buffered saline (PBS; Invitrogen) and incubated in calcium-free DMEM containing 4 mM EGTA (Millipore Sigma) and 10% low-calcium fetal bovine serum (FBS) for 10 minutes. This was followed by a switch to calcium-free DMEM supplemented with 2 mM CaCl₂, and cells were fixed at 6, 12, and 24 hours timepoints. Cells were fixed using ice-cold 1:1 methanol-acetone for 10 minutes. After fixation, cells were rinsed three times with PBS and blocked for 1 hour at room temperature using 1% bovine serum albumin (BSA). The following day, samples were incubated with primary antibodies (1:200 dilution) for 1 hour at room temperature, followed by incubation with secondary antibodies (1:500 dilution) for 30 minutes. Each antibody incubation step was followed by three PBS washes. Finally, samples were mounted using ProLong Diamond Antifade Mountant (Life Technologies) and stored in the dark at 4 °C until imaging.

Immunofluorescence images were acquired at room temperature using a Leica TCS SP8 STED 3X confocal microscope (Leica Microsystems) equipped with a 100× oil immersion objective (HC PL APO CS2). Excitation was provided by a white light laser spanning 470–670 nm, and depletion was performed using laser lines at 592, 660, and 775 nm. Image acquisition was carried out using LAS X software, followed by deconvolution with Huygens Pro (Scientific Volume Imaging). Post-processing was performed in ImageJ including cropping, resizing, grayscale inversion, brightness and contrast adjustments, and color lookup table modifications. Pixel resolution was set to approximately 25 nm × 25 nm.

### Immunofluorescence confocal microscopy

Cells were seeded on glass coverslips coated with collagen and cultured until they reached approximately 90% confluency. The cells were then fixed for 10 minutes at 4 °C using a chilled 1:1 acetone/methanol solution. Fixed samples were washed three times with PBS and blocked using a blocking buffer. Cells were incubated with a primary antibody (1:200 dilution) for 1 hour, followed by a secondary antibody (1:500 dilution) for 30 minutes at room temperature.

Imaging for border recruitment analysis was performed using a Leica Stellaris 5 microscope (Leica Microsystems) equipped with a 63× oil immersion objective (HC PL APO CS2). LAS X software was used for image acquisition, and ImageJ was employed for post-processing. For colocalization analysis, images were captured in a single z-plane and analyzed using the ‘Colo2’ module in ImageJ2, applying the Costes threshold regression method with a point spread function (PSF) setting of 3 and 50 Costes randomizations to calculate the Pearson correlation coefficient.

Confocal immunofluorescence images were further processed with grayscale inversion, pixel resizing, and contrast enhancement. A standardized image acquisition workflow was used across all samples to ensure consistency. Pixel resolution was set to approximately 20 nm × 20 nm.

### Molecular Dynamic (MD) simulation and analysis

Molecular dynamics (MD) simulations were conducted using GROMACS 2022.3 on the FARM high-performance computing cluster at the University of California, Davis, following previously described protocols^24,25,31^. Simulations were initiated using the OPLS-AA/L force field and the TIP4P water model, with a 10 Å cutoff applied for both van der Waals and electrostatic interactions. Electrostatic energies were computed using the particle mesh Ewald (PME) method with a grid spacing of 0.16 nm. Due to the lack of a crystal structure for the Dsg2 strand-swap heterodimer, we used the available crystal structures of the Dsg2 strand-swap homodimer (PDB: 5ERD), Pcad strand-swap homodimer (PDB: 4ZML), and Ecad strand-swap homodimer (PDB: 2O72). One Dsg2 monomer was replaced with either a Pcad or Ecad monomer by aligning the EC1 domain (residues 1–106) in PyMOL. The aligned structures were then prepared using PDBFixer for subsequent simulations.

Simulations began by placing the aligned structures at the center of a dodecahedral simulation box, ensuring a minimum distance of 1 nm from the box edges. The system was first subjected to energy minimization to eliminate unfavorable interactions, followed by equilibration under isothermal–isochoric and isothermal–isobaric conditions using a modified Berendsen thermostat and Berendsen barostat. The simulation box was solvated with water molecules and neutralized with ions, including 100 mM NaCl, 4 mM KCl, and 2 mM CaCl₂. After equilibration, a 50 ns molecular dynamics (MD) simulation was performed using a 2 fs integration time step. Structural stabilization was typically achieved by approximately 20 ns (Supplementary Fig. 3).

For Trp2 stabilization analysis, the gmx pairdist module was used to measure the distance between the center of mass (COM) of Trp2 from either Pcad or Ecad and the COM of the hydrophobic pocket from Dsg2. To measure the angle between the pre-hinge and post-hinge strands, their principal axes were determined using the gmx principal module. The angle was then calculated using the equation: 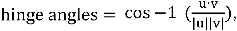 in which u and v are the a1 principal axis vectors.

### Statistical analysis

Statistical analyses were conducted using Python and Excel. As the study primarily involved pairwise comparisons, data were evaluated using two-sided Student’s *t*-tests. Corresponding *p*-values are reported in the figure panels.

## Acknowledgments

This research was supported by the National Institute of General Medical Sciences of the National Institutes of Health (R01GM121885 and R01GM133880). STED imaging was performed at the UC Davis Advanced Imaging Facility. FACS was performed at the Flow Cytometry Shared Resource, funded by the UC Davis Comprehensive Cancer Center Support Grant (CCSG) awarded by the National Cancer Institute (NCI P30CA093373).

## Data availability

All data are made available in the manuscript and supporting information.

## Reference

1 Gumbiner, B. M. Cell adhesion: the molecular basis of tissue architecture and morphogenesis. Cell 84, 345–357 (1996). 10.1016/s0092-8674(00)81279-9

2 Koch, P. J. et al. Targeted disruption of the pemphigus vulgaris antigen (desmoglein 3) gene in mice causes loss of keratinocyte cell adhesion with a phenotype similar to pemphigus vulgaris. J Cell Biol 137, 1091–1102 (1997). 10.1083/jcb.137.5.1091

3 Green, K. J. & Simpson, C. L. Desmosomes: new perspectives on a classic. J Invest Dermatol 127, 2499–2515 (2007). 10.1038/sj.jid.5701015

4 Michels, C., Buchta, T., Bloch, W., Krieg, T. & Niessen, C. M. Classical cadherins regulate desmosome formation. J Invest Dermatol 129, 2072–2075 (2009). 10.1038/jid.2009.17

5 Amagai, M. et al. Delayed assembly of desmosomes in keratinocytes with disrupted classic-cadherin-mediated cell adhesion by a dominant negative mutant. J Invest Dermatol 104, 27–32 (1995). 10.1111/1523-1747.ep12613462

6 Hatzfeld, M., Keil, R. & Magin, T. M. Desmosomes and Intermediate Filaments: Their Consequences for Tissue Mechanics. Cold Spring Harb Perspect Biol 9 (2017). 10.1101/cshperspect.a029157

7 Zimmer, S. E. & Kowalczyk, A. P. The desmosome as a dynamic membrane domain. Curr Opin Cell Biol 90, 102403 (2024). 10.1016/j.ceb.2024.102403

8 Shafraz, O. et al. E-cadherin binds to desmoglein to facilitate desmosome assembly. Elife 7 (2018). 10.7554/eLife.37629

9 Lewis, J. E., Jensen, P. J. & Wheelock, M. J. Cadherin function is required for human keratinocytes to assemble desmosomes and stratify in response to calcium. J Invest Dermatol 102, 870–877 (1994). 10.1111/1523-1747.ep12382690

10 Lenox, J. M. et al. Postnatal lethality of P-cadherin/desmoglein 3 double knockout mice: demonstration of a cooperative effect of these cell adhesion molecules in tissue homeostasis of stratified squamous epithelia. J Invest Dermatol 114, 948–952 (2000). 10.1046/j.1523-1747.2000.00976.x

11 Jones, J. C. Characterization of a 125K glycoprotein associated with bovine epithelial desmosomes. J Cell Sci 89 **(Pt** **2**), 207–216 (1988). 10.1242/jcs.89.2.207

12 Harrison, O. J. et al. The extracellular architecture of adherens junctions revealed by crystal structures of type I cadherins. Structure 19, 244–256 (2011). 10.1016/j.str.2010.11.016

13 Harrison, O. J. et al. Two-step adhesive binding by classical cadherins. Nat Struct Mol Biol 17, 348–357 (2010). 10.1038/nsmb.1784

14 Zhang, Y., Sivasankar, S., Nelson, W. J. & Chu, S. Resolving cadherin interactions and binding cooperativity at the single-molecule level. Proc Natl Acad Sci U S A 106, 109–114 (2009). 10.1073/pnas.0811350106

15 Parisini, E., Higgins, J. M., Liu, J. H., Brenner, M. B. & Wang, J. H. The crystal structure of human E-cadherin domains 1 and 2, and comparison with other cadherins in the context of adhesion mechanism. J Mol Biol 373, 401–411 (2007). 10.1016/j.jmb.2007.08.011

16 Boggon, T. J. et al. C-cadherin ectodomain structure and implications for cell adhesion mechanisms. Science 296, 1308–1313 (2002). 10.1126/science.1071559

17 Harrison, O. J. et al. Structural basis of adhesive binding by desmocollins and desmogleins. Proc Natl Acad Sci U S A 113, 7160–7165 (2016). 10.1073/pnas.1606272113

18 Troyanovsky, R. B. et al. Sorting of cadherin-catenin-associated proteins into individual clusters. Proc Natl Acad Sci U S A 118 (2021). 10.1073/pnas.2105550118

19 Stahley, S. N., Bartle, E. I., Atkinson, C. E., Kowalczyk, A. P. & Mattheyses, A. L. Molecular organization of the desmosome as revealed by direct stochastic optical reconstruction microscopy. J Cell Sci 129, 2897–2904 (2016). 10.1242/jcs.185785

20 Stahley, S. N. et al. Super-Resolution Microscopy Reveals Altered Desmosomal Protein Organization in Tissue from Patients with Pemphigus Vulgaris. J Invest Dermatol 136, 59–66 (2016). 10.1038/JID.2015.353

21 Dong, Y., Elgerbi, A., Xie, B., Choy, J. S. & Sivasankar, S. Actomyosin forces trigger a conformational change in desmoplakin within desmosomes. bioRxiv (2025). 10.1101/2024.11.19.624364

22 Dean, W. F. et al. Dsg2 ectodomain organization increases throughout desmosome assembly. Cell Adh Migr 18, 1–13 (2024). 10.1080/19336918.2024.2333366

23 Kudo, S., Caaveiro, J. M. & Tsumoto, K. Adhesive Dimerization of Human P-Cadherin Catalyzed by a Chaperone-like Mechanism. Structure 24, 1523–1536 (2016). 10.1016/j.str.2016.07.002

24 Xie, B. et al. Molecular mechanism for strengthening E-cadherin adhesion using a monoclonal antibody. Proc Natl Acad Sci U S A 119, e2204473119 (2022). 10.1073/pnas.2204473119

25 Xie, B., Xu, S., Schecterson, L., Gumbiner, B. M. & Sivasankar, S. Strengthening E-cadherin adhesion via antibody-mediated binding stabilization. Structure 32, 217–227 e213 (2024). 10.1016/j.str.2023.11.002

26 Vendome, J. et al. Molecular design principles underlying beta-strand swapping in the adhesive dimerization of cadherins. Nat Struct Mol Biol 18, 693–700 (2011). 10.1038/nsmb.2051

27 Al-Amoudi, A., Diez, D. C., Betts, M. J. & Frangakis, A. S. The molecular architecture of cadherins in native epidermal desmosomes. Nature 450, 832– 837 (2007). 10.1038/nature05994

28 Schinner, C. et al. Stabilization of desmoglein-2 binding rescues arrhythmia in arrhythmogenic cardiomyopathy. JCI Insight 5 (2020). 10.1172/jci.insight.130141

29 Schmitt, C. J. et al. Homo- and heterotypic cell contacts in malignant melanoma cells and desmoglein 2 as a novel solitary surface glycoprotein. J Invest Dermatol 127, 2191–2206 (2007). 10.1038/sj.jid.5700849

30 Koirala, R. et al. Inside-out regulation of E-cadherin conformation and adhesion. Proc Natl Acad Sci U S A 118 (2021). 10.1073/pnas.2104090118

31 Xie, B., Xu, S. & Sivasankar, S. Outside-in engineering of cadherin endocytosis using a conformation strengthening antibody. Nat Commun 16, 1157 (2025). 10.1038/s41467-025-56478-6

32 Simcha, I. et al. Differential nuclear translocation and transactivation potential of beta-catenin and plakoglobin. J Cell Biol 141, 1433–1448 (1998). 10.1083/jcb.141.6.1433

33 Huber, A. H. & Weis, W. I. The structure of the beta-catenin/E-cadherin complex and the molecular basis of diverse ligand recognition by beta-catenin. Cell 105, 391–402 (2001). 10.1016/s0092-8674(01)00330-0

34 Lewis, J. E. et al. Cross-talk between adherens junctions and desmosomes depends on plakoglobin. J Cell Biol 136, 919–934 (1997). 10.1083/jcb.136.4.919

35 Priest, A. V., Koirala, R. & Sivasankar, S. Cadherins can dimerize via asymmetric interactions. FEBS Lett 596, 1639–1646 (2022). 10.1002/1873-3468.14373

36 Kurzen, H., Munzing, I. & Hartschuh, W. Expression of desmosomal proteins in squamous cell carcinomas of the skin. J Cutan Pathol 30, 621–630 (2003). 10.1034/j.1600-0560.2003.00122.x

37 Jurcic, V., Kukovic, J. & Zidar, N. Expression of desmosomal proteins in acantholytic squamous cell carcinoma of the skin. Histol Histopathol 30, 945– 953 (2015). 10.14670/HH-11-599

38 Chidgey, M. & Dawson, C. Desmosomes: a role in cancer? Br J Cancer 96, 1783–1787 (2007). 10.1038/sj.bjc.6603808

39 Yashiro, M., Nishioka, N. & Hirakawa, K. Decreased expression of the adhesion molecule desmoglein-2 is associated with diffuse-type gastric carcinoma. Eur J Cancer 42, 2397–2403 (2006). 10.1016/j.ejca.2006.03.024

40 Shelton, W. T. et al. Desmoglein-2 harnesses a PDZ-GEF2/Rap1 signaling axis to control cell spreading and focal adhesions independent of cell-cell adhesion. Sci Rep 11, 13295 (2021). 10.1038/s41598-021-92675-1

41 Roberts, B. J. et al. Palmitoylation of Desmoglein 2 Is a Regulator of Assembly Dynamics and Protein Turnover. J Biol Chem 291, 24857–24865 (2016). 10.1074/jbc.M116.739458

42 Samuelov, L., Sprecher, E. & Paus, R. The role of P-cadherin in skin biology and skin pathology: lessons from the hair follicle. Cell Tissue Res 360, 761– 771 (2015). 10.1007/s00441-015-2114-y

43 Vieira, A. F. & Paredes, J. P-cadherin and the journey to cancer metastasis. Mol Cancer 14, 178 (2015). 10.1186/s12943-015-0448-4

