## Supplemental Information for "P-cadherin and desmoglein-2 interact as strand-swap dimers and facilitate desmosome assembly"

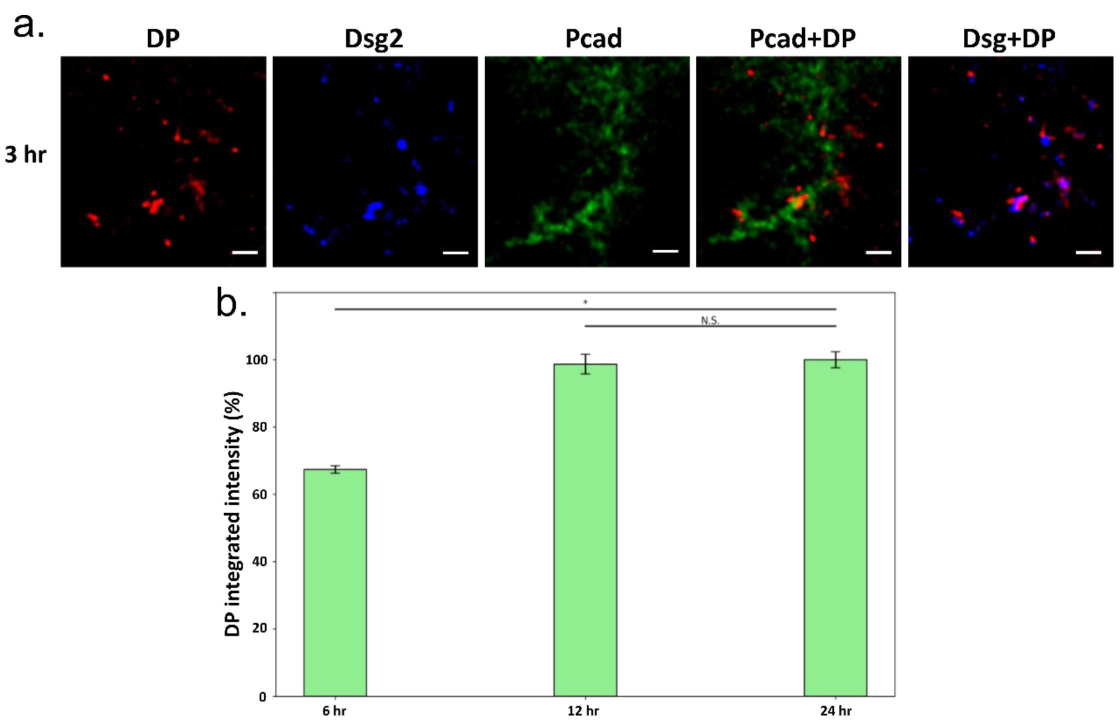


**Supplementary Figure 1. STED analysis of DP intensity in WT-Pcad cells at cell-cell contacts, measured at different timepoints following calcium switch. (a)** Representative image of cell-cell junction, fixed 3hr after calcium switch. DP expression was relatively scattered, and a consistent “railroad track” structure was not observed. **(b)** Quantification of DP intensity at cell-cell contact at 6 hr, 12 hr and 24 hr timepoints. At 6 hours, the integrated DP intensity was relatively low in DP railroad tracks. Beyond 12 hours, the DP intensity increased and stabilized due to desmosome maturation. Statistical analyses were performed between 24 hours samples and other samples. For 6-hour samples, the P value is 3.73E-02. For 12-hour samples, the P value is 0.950. Measurements are from three independent replicates and n is 43, 30, and 36 for 6-hour, 12-hour, and 24-hour samples, respectively.

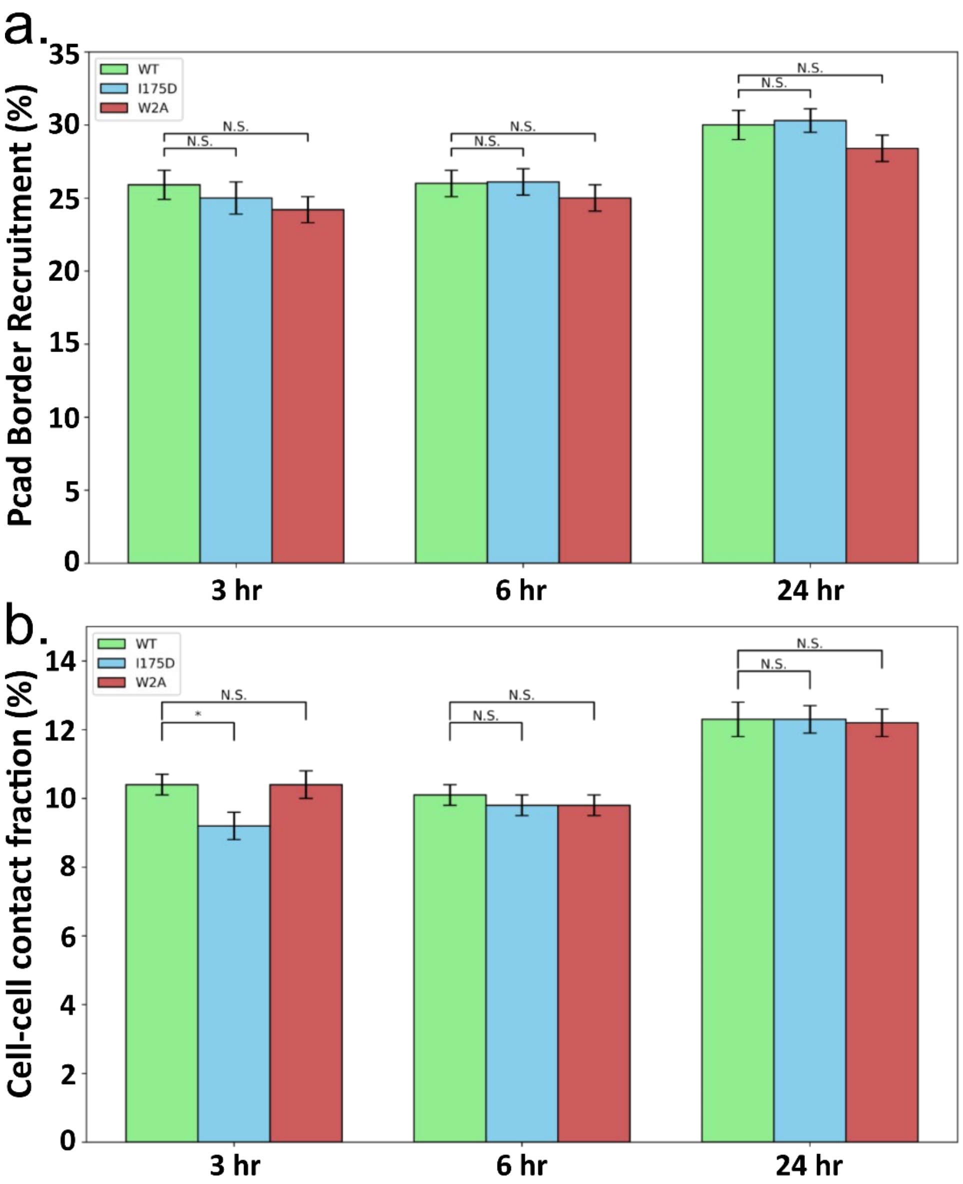


**Supplementary Figure 2. P-cadherin membrane recruitment level at different time points following calcium switch in cells expressing WT-Pcad, I175D-Pcad, and W2A-Pcad. (a)** Quantification of Pcads localized at cell-cell contacts. WT (green), I175D (blue), and W2A (red). **(b)** Fraction of cell border that is engaged in cell-cell contact. These results suggest that Pcad expression at cell-cell contacts and the fraction of cell-cell contact are roughly comparable between different cells. Statistical analyses were performed between WT-Pcad samples and other mutant Pcad samples. In Pcad recruitment analysis, for 3-hour samples, the P value for I175D-Pcad is 0.544 and P value for W2A-Pcad is 0.190. For 6-hour samples, the P value for I175D-Pcad is 0.963 and P value for W2A-Pcad is 0.425. For 24-hour samples, the P value for I175D-Pcad is 0.396 and P value for W2A-Pcad is 0.229. In cell-cell contact fraction analysis, for 3-hour samples the P value for I175D-Pcad is 1.70E-02 and P value for W2A-Pcad is 0.990. For 6-hour samples, the P value for I175D-Pcad is 0.457 and P value for W2A-Pcad is 0.494. For 24-hour samples, the P value for I175D-Pcad is 0.988 and P value for W2A-Pcad is 0.861. These measurements were calculated using the same dataset from Figure 3d and 3e.

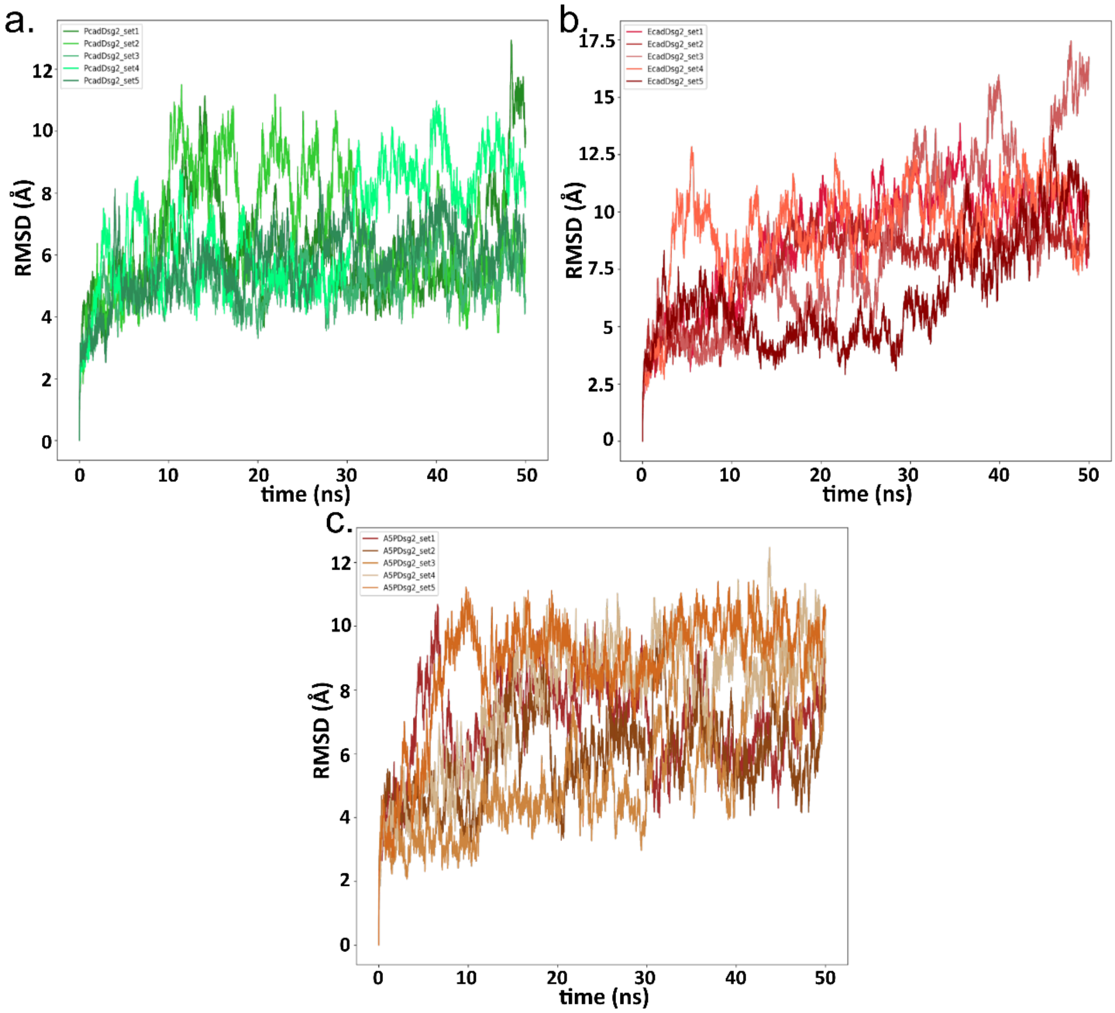


**Supplementary Figure 3. Protein backbone Root Mean Square Deviation (RMSD) during MD simulations, calculated relative to the initial structures at the start of simulation.** RMSD values measured for every set in three different initial structures: **(a)** WT-Pcad:WT-Dsg2, **(b)** WT-Ecad:WT-Dsg2, and **(c)** A5P-Pcad:WT-Dsg2. While the Pcad:Dsg2 structures had lower RMSD values, the Ecad:Dsg2 structures showed greater RMSD values throughout the simulations.

a. b.

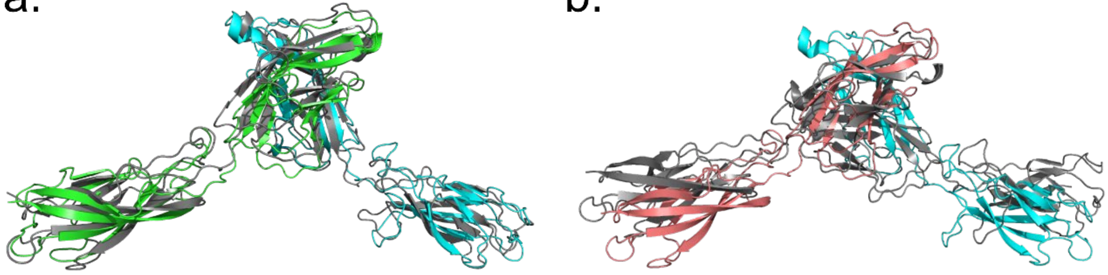

**Supplementary Figure 4. Protein structure comparison between structures at the beginning and end of the MD simulation. (a)** Representative WT-Pcad:WT-Dsg2 initial structure aligned with final structure shows preservation of S-dimer conformation. The colored structure (Pcad in green and Dsg2 in cyan) is the initial structure and the grey structure is the final structure after MD. **(b)** Representative WT-Ecad:WT-Dsg2 initial structure aligned with final structure reveals a visible deformation from initial S-dimer structure. The colored structure (Ecad in red and Dsg2 in cyan) is the initial structure, and the grey structure is the final structure after MD.

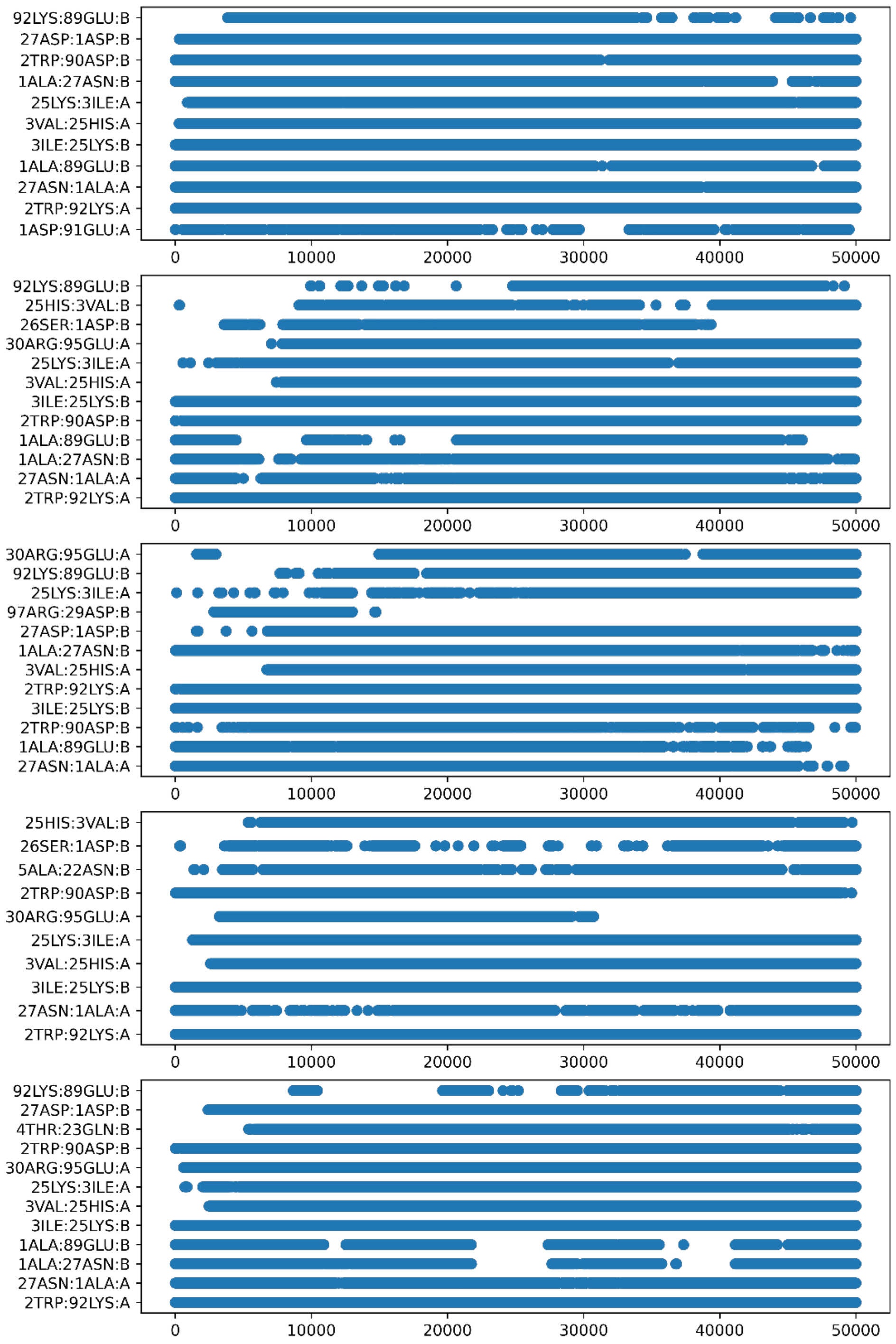


**Supplementary Figure 5. Persistent electrostatic interactions between WT-Pcad:WT-Dsg2 S-dimers (5 sets included).** For each of the simulation sets, electrostatic interactions lasting for more than 30% of total frames were considered as persistent.

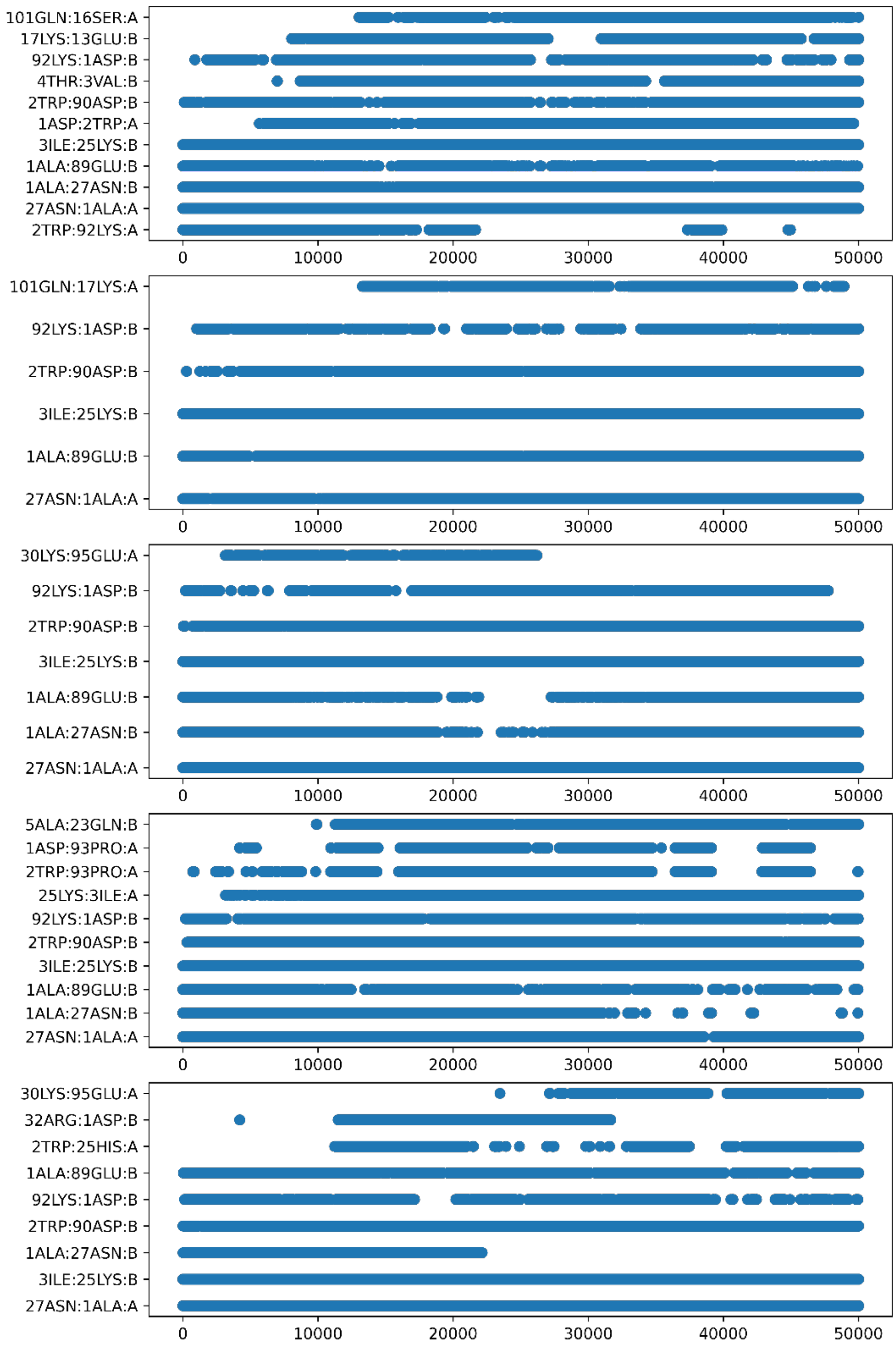


**Supplementary Figure 6. Persistent electrostatic interactions between WT-Ecad:WT-Dsg2 S-dimers (5 sets included).** For each of the simulation sets, electrostatic interactions lasting for more than 30% of total frames were considered as persistent. There are fewer persistent electrostatic interactions compared to WT-Pcad:WT-Dsg2 S-dimers.

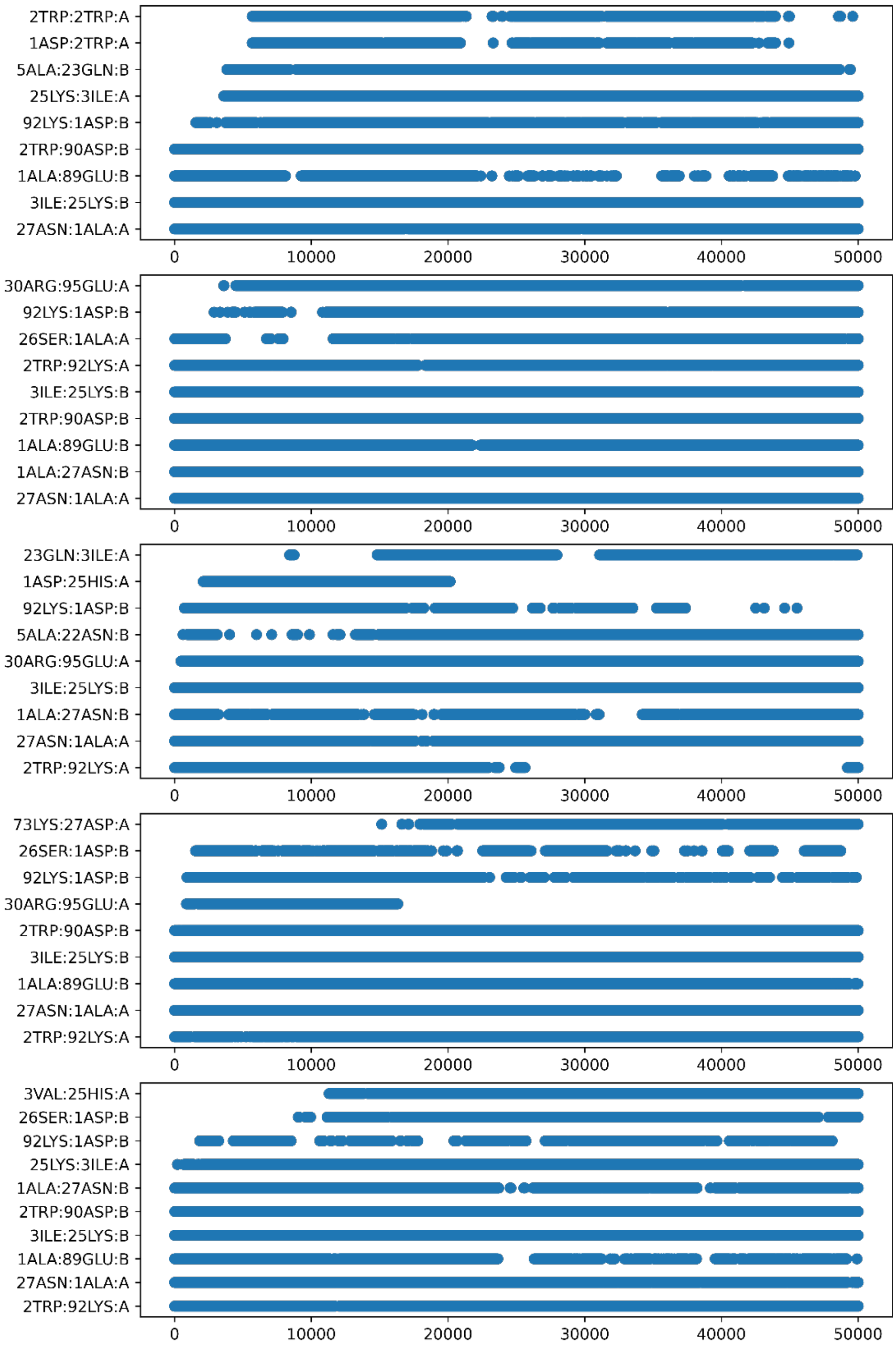


**Supplementary Figure 7. Persistent electrostatic interactions between A5P-Pcad:WT-Dsg2 S-dimers (5 sets included).** For each of the simulation sets, electrostatic interactions lasting for more than 30% of total frames were considered as persistent. There are fewer persistent electrostatic interactions compared to WT-Pcad:WT-Dsg2 S-dimers.

a. Desmosomal cadherins

| | 1 | 2 | 3 | 4 | 5 | 6 | 7 | 8 | 9 | 10 | 11 | 12 |
|---|---|---|---|---|---|---|---|---|---|---|---|---|
| Dsg1 | E | W | I | K | F | A | A | A | C | R | E | G |
| Dsg3 | E | W | V | K | F | A | K | P | C | R | E | G |
| Dsg4 | E | W | I | K | F | A | A | A | C | R | E | G |
| Dsc1 | R | W | A | P | I | P | A | S | L | M | E | N |
| Dsc2 | R | W | A | P | I | P | C | S | M | Q | E | N |
| Dsc3 | R | W | A | P | I | P | C | S | M | L | E | N |

b. Classical cadherins

| | 1 | 2 | 3 | 4 | 5 | 6 | 7 | 8 | 9 | 10 | 11 | 12 |
|---|---|---|---|---|---|---|---|---|---|---|---|---|
| M-Ecad | D | W | V | I | P | P | I | S | C | P | E | N |
| M-Pcad | E | W | V | M | P | P | I | F | V | P | E | N |
| M-Ncad | D | W | V | I | P | P | I | N | L | P | E | N |
| M-Rcad | D | W | V | I | P | P | I | N | V | P | E | N |
| C-Ecad | D | W | V | I | P | P | I | S | C | P | E | N |
| C-Pcad | D | W | V | V | A | P | I | S | V | P | E | N |
| C-Ncad | D | W | V | I | P | P | I | N | L | P | E | N |
| C-Rcad | D | W | V | I | P | P | I | N | V | P | E | N |

**Supplementary Figure 8. β strand alignment for different classical and desmosomal cadherins.** Sequence alignment over β strand across **(a)** human desmosomal cadherins and **(b)** classical cadherin from mouse and chimpanzee. This result suggests that Ala5-Pro6 hinge is relatively restricted only to Dsg2 and primate species Pcad.

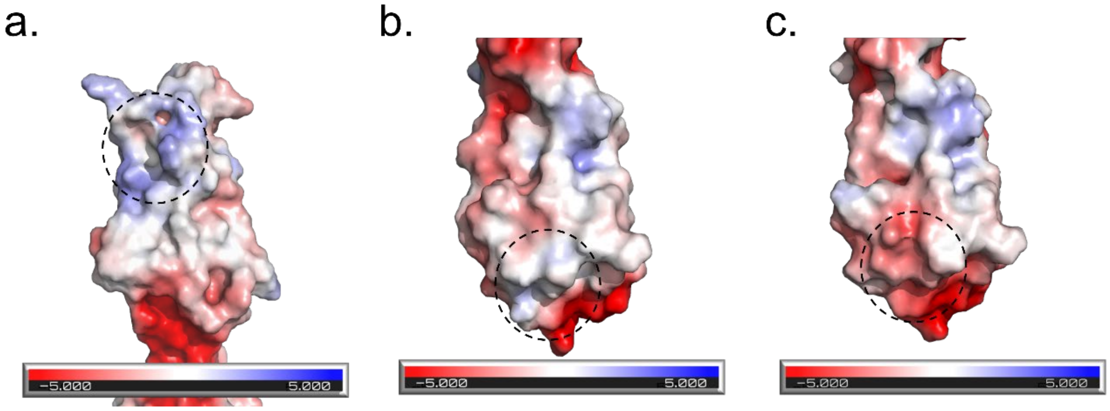


**Supplementary Figure 9. Electrostatic surface potential analysis for putative cis interface of Dsg2, Pcad and Ecad. (a)** Putative cis interface of Dsg2 EC1 domain (marked region) shows positively charged surface. **(b)** cis interface of Pcad EC1 domain (marked region) also shows a slight positive charge. **(c)** cis interface of Ecad EC1 domain (marked region) shows negatively charged surface which promotes interaction with Dsg2 in a cis orientation.
